# Comprehensive phylogenomics of *Methylobacterium* reveals four evolutionary distinct groups and underappreciated phyllosphere diversity

**DOI:** 10.1101/2022.03.12.484109

**Authors:** Jean-Baptiste Leducq, David Sneddon, Malia Santos, Domitille Condrain-Morel, Geneviève Bourret, N. Cecilia Martinez-Gomez, Jessica A. Lee, James A. Foster, Sergey Stolyar, B. Jesse Shapiro, Steven W. Kembel, Jack M Sullivan, Christopher J. Marx

**Affiliations:** University of Idaho; Université du Québec à Montréal; UC, Berkeley; NASA Ames Research Center; Université de Montréal; McGill University

**Keywords:** *Methylobacterium*, *Methylorubrum*, species concept in bacteria, horizontal gene transfers, genome architecture, core genome, lineage tree, species tree, phyllosphere

## Abstract

*Methylobacterium* is a group of methylotrophic microbes associated with soil, fresh water, and particularly the phyllosphere, the aerial part of plants that has been well-studied in terms of physiology but whose evolutionary history and taxonomy are unclear. Recent work has suggested that *Methylobacterium* is much more diverse than thought previously, questioning its status as an ecologically and phylogenetically coherent taxonomic genus. However, taxonomic and evolutionary studies of *Methylobacterium* have mostly been restricted to model species, often isolated from habitats other than the phyllosphere, and have yet to utilize comprehensive phylogenomic methods to examine gene trees, gene content, or synteny. By analyzing 189 *Methylobacterium* genomes from a wide range of habitats, including the phyllosphere, we inferred a robust phylogenetic tree while explicitly accounting for the impact of horizontal gene transfers. We showed that *Methylobacterium* contains four evolutionary distinct groups of bacteria (namely A, B, C, D), characterized by different genome size, GC content, gene content and genome architecture, revealing the dynamic nature of *Methylobacterium* genomes. In addition of recovering 59 described species, we identified 45 candidate species, mostly phyllosphere-associated, stressing the significance of plants as a reservoir of *Methylobacterium* diversity. We inferred an ancient transition from a free-living lifestyle to association with plant roots in *Methylobacteriaceae* ancestor, followed by phyllosphere association of three of the major groups (A, B, D), which early branching in *Methylobacterium* history was heavily obscured by HGT. Together, our work lays the foundations for a thorough redefinition of *Methylobacterium* taxonomy, beginning with the abandon of *Methylorubrum*.

## Introduction

For billions of years, bacteria evolved rapidly through vertical and horizontal gene transmission, mutation, selection, diversification, and extinction. These evolutionary processes allowed bacteria to conquer every biome and living host on Earth and, at the same time, resulted in blurring most traces of their ancient history (Louca *et al*., 2018). In the past thousands of years, humans have increasingly imposed new selective pressures on bacterial evolution, through bacterial host domestication and ecosystem perturbations (Gillings and Paulsen, 2014). Ironically, the human perception of microbial life was until recently limited to the diversity we could “see” (through cultivation) and “use” (through domestication), representing only an infinitesimal proportion of the bacterial diversity in nature (Hugenholtz, 2002). As a result, bacterial diversity, evolution and speciation concepts remain fuzzy and largely biased (Shapiro *et al*., 2016). Yet, the advent of high-throughput sequencing technologies, and our awakening to the essential role of bacteria in every living system has spurred research into the evolutionary processes shaping the microbial world (Koonin *et al*., 2021).

*Methylobacterium* (*Alphaproteobacteria, Rhizobiales, Methylobacteriaceae*) is a well-studied group of bacteria that are abundant and widespread in every plant microbiome (Corpe and Rheem, 1989; Keppler *et al*., 2006). Easy to isolate and to cultivate, thanks to a pink coloration due to carotenoids and their ability to use methanol as sole carbon source (Clarke, 1983; Anthony, 1991; Keppler *et al*., 2006), methylobacteria are also essential players in plant functions, like growth stimulation (Ivanova *et al*., 2001; Madhaiyan *et al*., 2005, 2007), heavy metal sequestration (Madhaiyan *et al*., 2007), protection against phytopathogens and nitrogen fixation (Dourado *et al*., 2015), sparking increasing interest in their use in plant biotechnology applications (Ryu *et al*., 2006; Lee *et al*., 2006; Madhaiyan *et al*., 2007).

Recently, Green and Ardley (2018) questioned the taxonomy of *Methylobacterium*, noticing a *“greater degree of phenotypic and genotypic heterogeneity than would normally be expected for a single genus*.” Accordingly, these authors proposed to split the genus in three distinct taxa corresponding to monophyletic groups in the 16S rRNA ribosomal gene phylogeny (groups A, B and C). Group A, containing the *Methylobacterium* type species *M. organophilum*, was retained as *Methylobacterium*. For group B, which included the model species *M. extorquens*, the authors proposed a new genus: *Methylorubrum*. Finally, the authors suggested that group C, including *M. aquaticum* and *M. nodulans*, should constitute a distinct genus, pending future genetic and phenotypic investigations. The *Methylobacterium* reclassification has been pointed out as problematic, because of the low phylogenetic resolution of the 16S rRNA gene, and because no genus name was proposed for strains that were not retained in *Methylorubrum* or *Methylobacterium*, which could potentially render either new genus as paraphyletic (Hördt *et al*., 2020; Leducq *et al*., 2022). Accordingly, the taxonomy of *Methylobacterium* was reexamined by coupling genome-wide DNA-DNA hybridization and phenotypic information for 63 strains, each representative of a described species (Alessa *et al*., 2021). Alessa *et al*. confirmed Green and Ardley’s (2018) observation that group C was phenotypically and genetically distinct from other groups, but they also showed that *Methylorubrum* (group B) was embedded within *Methylobacterium* (group A), forming a homogeneous group, and proposed to merge *Methylobacterium* and *Methylorubrum* back into a single genus.

The evolutionary history of *Methylobacterium* remains poorly resolved for several reasons. First, phylogenetic relationships among and within groups are often inconsistent depending upon the chosen marker gene (Green and Ardley, 2018; Leducq *et al*., 2022). Such inconsistent phylogenetic signals suggest that these marker genes had different evolutionary histories, perhaps due to horizontal gene transfer (HGT) or incomplete lineage sorting (ILS), illustrating the dynamic nature of bacterial genome evolution and the limitations of bacterial taxonomy based on a limited number of gene phylogenies (Castillo-Ramírez and González, 2008; Creevey *et al*., 2011). Second, Alessa *et al*. (2021) based their *Methylobacterium* taxonomy on DNA-DNA hybridization methods, which are widely used to classify prokaryotic species, but are not phylogenetic methods *per se*, as they do not account for ancestry. They also validated their taxonomy using a phylogenetic tree based on concatenated protein sequences of core genes but did not present evaluations of the uncertainty in the resulting tree. Finally, phylogenies based on concatenated gene alignments assume the same tree for each gene, and thus do not take into account potential ILS and HGT affecting topology and branch lengths differentially in each individual gene trees. With the onset of genomics in evolutionary studies, several coalescent methods have been developed to reconstruct the phylogeny and solve the taxonomy of organisms with complex evolutionary history like bacteria (Davidson *et al*., 2015). For instance, coalescent-based phylogenetic methods like Astral (Mirarab *et al*., 2014) and SVDquartets (Chifman and Kubatko, 2014) allow the reconstruction of a consensus tree (the lineage tree) taking into account different levels of ILS and HGT among individual gene trees.

Although more than 60 *Methylobacterium* species have been described so far (Green and Ardley, 2018; Chen *et al*., 2019; Feng *et al*., 2020; Jia *et al*., 2020; Kim, Chhetri, Kim, Lee, *et al*., 2020; Kim, Chhetri, Kim, Kim, *et al*., 2020; Ten *et al*., 2020; Jiang *et al*., 2020; Pascual *et al*., 2020; Alessa *et al*., 2021), available genomic and phenotypic information was until recently limited to a few model species, mostly from groups B and C, and mostly isolated from anthropogenically impacted environments, and in rare cases from plants (Marx *et al*., 2012; Tani *et al*., 2015; Minami *et al*., 2016; Morohoshi and Ikeda, 2016; Belkhelfa *et al*., 2018). Surveys of *Methylobacterium* diversity associated with plants have mainly focused on the rhizosphere, especially in crop species (Sy *et al*., 2001; Jourand *et al*., 2004; Grossi *et al*., 2020). Recent studies however revealed that the phyllosphere of model plant species like *A. thaliana* (Helfrich *et al*., 2018), of wheat (Zervas *et al*., 2019), and of natural temperate forests (Leducq *et al*., 2022) are major reservoirs of undescribed *Methylobacterium* diversity, most of which belongs to group A (Leducq *et al*., 2022).

Here, we explored *Methylobacterium* diversity from an evolutionary genomic perspective. We *de novo* annotated 189 *Methylobacterium* genomes, including 62 strains isolated from temperate forest, wheat, and *Arabidopsis* phyllosphere, and 127 additional genomes that represent the remainder of the *Methylobacterium* species described so far. Using different phylogenomic approaches, we reconstructed the *Methylobacterium* evolutionary tree from 384 *Methylobacteriaceae* core genes and showed that the genus is consistently constituted of four monophyletic groups: A, B, C and D. Gene content and especially the highly dynamic core genome architecture predicted the four *Methylobacterium* groups remarkably well. We estimated that *Methylobacterium* includes at least 104 species, of which only 59 were previously described. Most of the undescribed species were assigned to groups A and D and were isolated from plant leaves, stressing the significance of the phyllosphere as a reservoir of *Methylobacterium* diversity. Our inferences of the *Methylobacterium* evolutionary tree also suggest an ancient transition from a free-living lifestyle to association with plant roots in *Methylobacteriaceae* ancestor, followed by phyllosphere association of three of the major groups (A, B, D), which early branching in *Methylobacterium* history was heavily obscured by HGT. Finally, our comprehensive phylogenetic analysis of *Methylobacterium* lays the foundation for a profound redefinition of its taxonomy, beginning with the abandon of *Methylorubrum*.

## Results

### Definition *of the* Methylobacteriaceae *core genome*

We assembled a collection of 213 *Methylobacteriaceae* genomes, including 189 *Methylobacterium* and 24 genomes from related genera as outgroups (*Microvirga*: n=22; *Enterovirga*: n=2). Most *Methylobacterium* (*n*=98) and all outgroup genomes (n=24) came from distinct studies (Dataset S1). We included 29 genomes from *Methylobacterium* type strains recently sequenced (Alessa *et al*., 2021; Bijlani *et al*., 2021), hence covering most *Methylobacterium* species described so far. We also included 38 genomes available from two large surveys of the *Arabidopsis* and wheat phyllospheres (Helfrich *et al*., 2018; Zervas *et al*., 2019), and sequenced 24 additional genomes of isolates from a large survey of the temperate forest phyllosphere (Leducq *et al*., 2022), hence extending our dataset to the leaf-associated *Methylobacterium* diversity. The 24 newly assembled genomes had 41 to 405 scaffolds (depth: 188-304x) for a total size (5-7Mb) and average GC content (67-70%) in the expected range for *Methylobacterium* genomes (Dataset S2). We annotated 184 genomes *de novo*, excluding 29 genomes that were not published at the time of the analysis (Alessa *et al*., 2021; Bijlani *et al*., 2021) through the same pipeline (RAST) and after excluding hypothetical proteins, repeat and mobile elements, we identified 9,970 unique gene annotations (i.e., regardless of copy number: Dataset S3), with on average 2637 (SD: 210) unique gene annotations per genome. We identified 893 candidate core genes, i.e., genes that were present in a single copy in at least 90% of *Methylobacteriaceae* genomes. After filtering for missing data and false duplications attributable to large variations among genome assembly qualities (Figures S1, S2), we identified 384 *Methylobacteriaceae* core genes (Dataset S4) for which the complete nucleotide sequences could be retrieved for at least 181 out of 184 genomes. We repeated the RAST annotation for recently sequenced genomes from 29 *Methylobacterium* species type strains that were not available during our initial survey (Alessa *et al*., 2021; Bijlani *et al*., 2021). Doing this, we slightly extended the number of unique gene annotation in *Methylobacteriaceae* (*n* = 10,190). We confirmed that the 384 previously identified genes were part of the *Methylobacteriaceae* core genome and retrieved each core gene nucleotide sequence for at least 26 out of these 29 genomes. Our final dataset consisted of 213 genomes for which we retrieved 327 to 384 core genes nucleotide sequences (average, SD: 381 ± 6).

### *Inference of the* Methylobacteriaceae *lineage tree*

We reconstructed the lineage tree of *Methylobacteriaceae* from 213 genomes from the 384 core gene nucleotide sequences using three complementary approaches in order to assess the effect of ILS and HGT in the evolutionary history of *Methylobacterium*. First, we used RAxML to determine a maximum-likelihood tree (512 replicated tree; bootstraps) from concatenated alignments of the core gene nucleotide sequences, assuming 57 groups of genes (partitions) with different substitution models (partitions determined in IQ-tree; GTRCAT model of substitution), but the same evolutionary tree for all genes, hence not accounting for ILS or HGT (Figure 1a). Second, we used ASTRAL, a coalescent-based method combining Maximum-Likelihood (ML) trees determined for each core gene independently (RAxML, GTRGAMMA model, 1,000 replicated tree), accounting for ILS among genes (Figure 1b, Figure S3a). Third, we used SVDquartets, a coalescent-based method estimating the tree for each possible combination of four genomes and assuming all nucleotide sites are unlinked in the concatenated alignment of 384 genes, hence accounting for ILS and HGT both within and among genes (Figure 1c, Figure S3b). In all lineage trees rooted on *Microvirga* and *Enterovirga, Methylobacterium* was monophyletic and consisted of four groups of genomes, consistently monophyletic and strongly supported, regardless of the method used (nodal support: 100% in RAxML and SVDquartets trees; local posterior probability: 1.0 in the ASTRAL tree; Figure 1a,b,c). Group C always formed the most basal group of *Methylobacterium*, confirming previous observations (Green and Ardley, 2018; Alessa *et al*., 2021; Leducq *et al*., 2022). Group B regrouped clades B, formerly *Methylorubrum* (Green and Ardley, 2018) and B2 (Alessa *et al*. 2021; corresponding to clade A4 in Leducq *et al*. 2022). Most strains previously assigned to clade A (Green and Ardley, 2018) were distributed across two distinct monophyletic groups that we named A and D (Figure 1a). Group A included clades A2, A3, A4 and A5 described by Alessa *et al*. (2021) and corresponded to clades A5, A10, A19 and A7+A8 described by Leducq *et al*. (2022), respectively. Group D corresponded to clade A1 proposed by Alessa *et al*. (2021) and clades A1, A2 and A3 proposed by Leducq *et al*. (2022).

**Figure 1:**
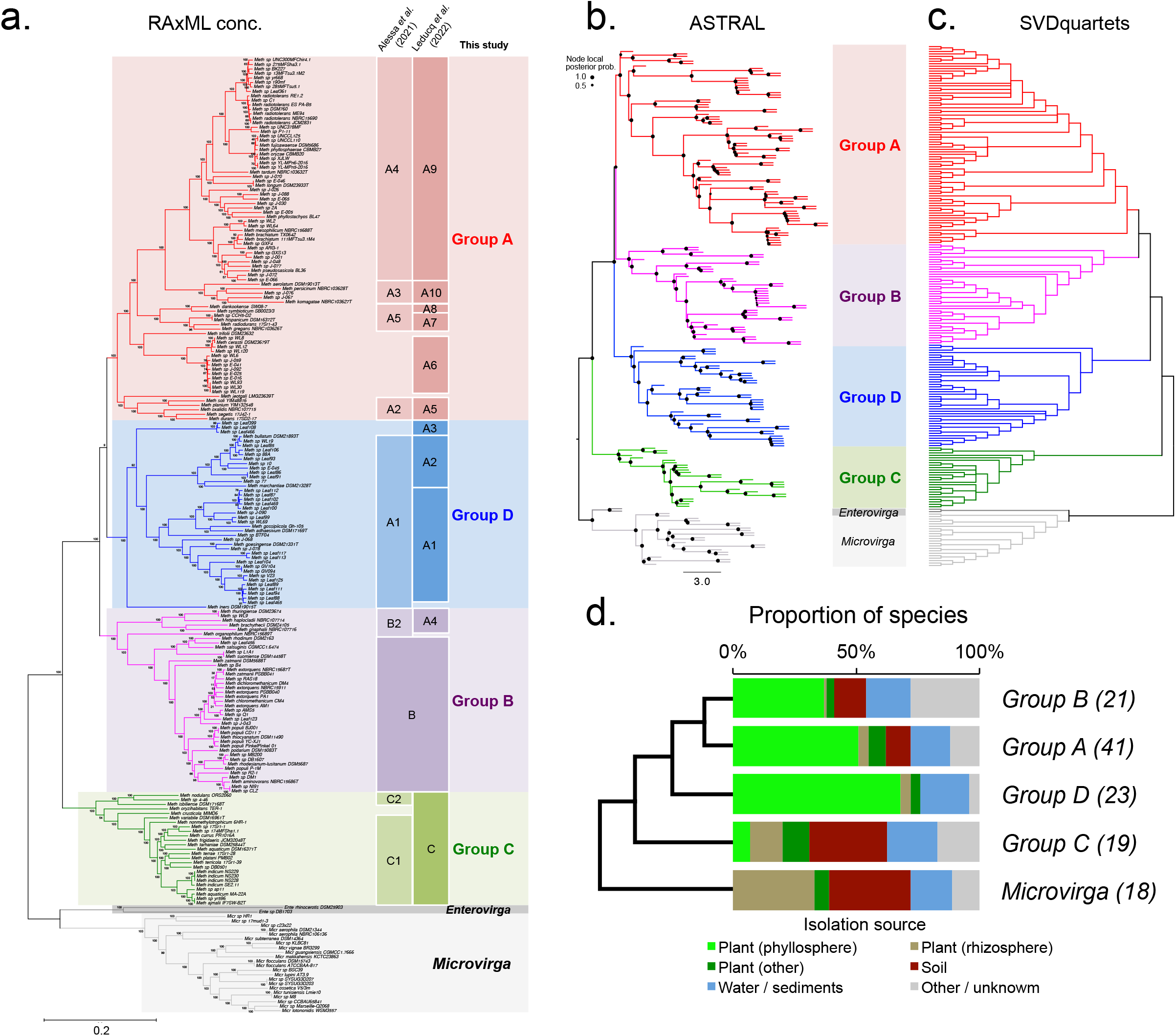
*Methylobacteriaceae* lineage trees inferred from 213 genomes. a) Best tree from RAxML ML search on the concatenated alignments of 384 core gene nucleotide sequences (GTRCAT model, 512 replicated trees), rooted on *Microvirga* and *Enterovirga* (grey). Colors indicate the four major *Methylobacterium* groups: A (red), B (purple), C (green) and D (blue). Correspondence with clades described by previous studies is indicated. b) ASTRAL tree inferred from 384 core gene ML trees. Each gene ML tree was inferred assuming a GTRgamma model (1,000 replicated trees; nodes with less than 10% of support collapsed) and combined in ASTRAL-III. Branch lengths are in coalescent units. Nodal support values represent local posterior probability. c) SVD quartet tree inferred from the concatenated alignments of 384 core gene nucleotide sequences. Nodes supported by less that 75% of quartets were collapsed. d) Main isolation sources of species from *Methylobacterium* group and *Microvirga* (see Table 1). For each group, ordered according to a consensus tree (see panels b and c), the number of species is indicated in parenthesis

Groups A, B and D consistently formed a monophyletic group (nodal support: 100% in RAxML and SVDquartets trees; local posterior probability: 1.0 in the ASTRAL tree); however, phylogenetic relationships among groups A, B and D were more challenging to assess. Groups A, B, and D could not be resolved with the RAxML tree (nodal support = 9%; Figure 1a). Group D was sister to groups A and B according to ASTRAL (local posterior probability: 0.8; Figure 1b) and SVDquartets trees (nodal support: 100%; Figure 1c). We evaluated differences between the three lineage tree topologies using the Robinson-Foulds (RF) distance metric in PAUP (Wilgenbusch and Swofford, 2003). RAxML and ASTRAL lineage tree topologies were more similar to each other (RF = 0.181) than with the SVDquartets tree (RF = 0.225 and 0.289, respectively; Figure S4). In order to determine whether the difference between the RAxML and other trees was higher than expected by chance, we estimated the distribution of RF distance between each replicate tree of the RAxML search for the lineage tree (512 replicates). The normalized RF value ranges from 0.028 to 0.113 (RF= 0.069±0.015), indicating that the differences observed between lineage trees were larger than expected by chance (Figure S4), and suggesting that ILS and HGT among core genes had a significant impact on the *Methylobacteriaceae* lineage tree. The larger difference between the SVDquartets tree and other trees also suggested that recombination within core genes also occurred during *Methylobacteriaceae* evolution, although without affecting the relationship among the four major groups (C/D/(A,B)).

### *Inference of the* Methylobacteriaceae *taxonomy and species tree*

We classified *Methylobacteriaceae* genomes into 124 species using a 97% threshold on percentage nucleotide similarity (*PNS*; analogous to average nucleotide identity; (Mende *et al*., 2013; Chun and Rainey); Dataset S5) on the core genome (concatenated alignments of 384 core genes; 361,403 bp). In the outgroups, we identified 2 *Enterovirga* species and 18 *Microvirga* species. We identified 104 *Methylobacterium* species (1 to 9 genomes per species), of which 59 included the type strain for at least one described species (Table 1; Dataset S5). *M. extorquens, M. chloromethanicum* and *M. dichloromethanicum* type strains were assigned to the same species (*PNS* range: 97.61-99.68%), as previously reported (Alessa *et al*., 2021). *M. populi* and *M. thiocyanatum* type strains were assigned to the same species (*PNS* range: 98.97%-99.08%), as previously reported (Alessa *et al*., 2021). *M. phyllosphaerae, M. ozyzae* and *M. fujisawaense* type strains were assigned to the same species (99.23-100%), as previously reported (Alessa *et al*., 2021). We identified 45 candidate species that included no type strain, and thus corresponded to new candidate *Methylobacterium* species (Table 1; Dataset S5). We numbered these candidate species from *M. sp*. 001 to *M. sp*. 045. We used the 124 identified species to infer the *Methylobacteriaceae* species trees with SVDquartets (Figure S5a) and ASTRAL (Figure S5b). Although the two species trees were not strictly identical (normalized RF distance = 0.234), the monophyly and relationships among the four main groups was consistent between ASTRAL and SVDquartets species trees (C/D/(AB); Figures S5a,b), and with ASTRAL and SVDquartets lineage trees (Figure 1b,c). Each group of genomes assigned to the same species was also monophyletic and strongly supported in lineage trees (Figure 1).

**Table 1:**
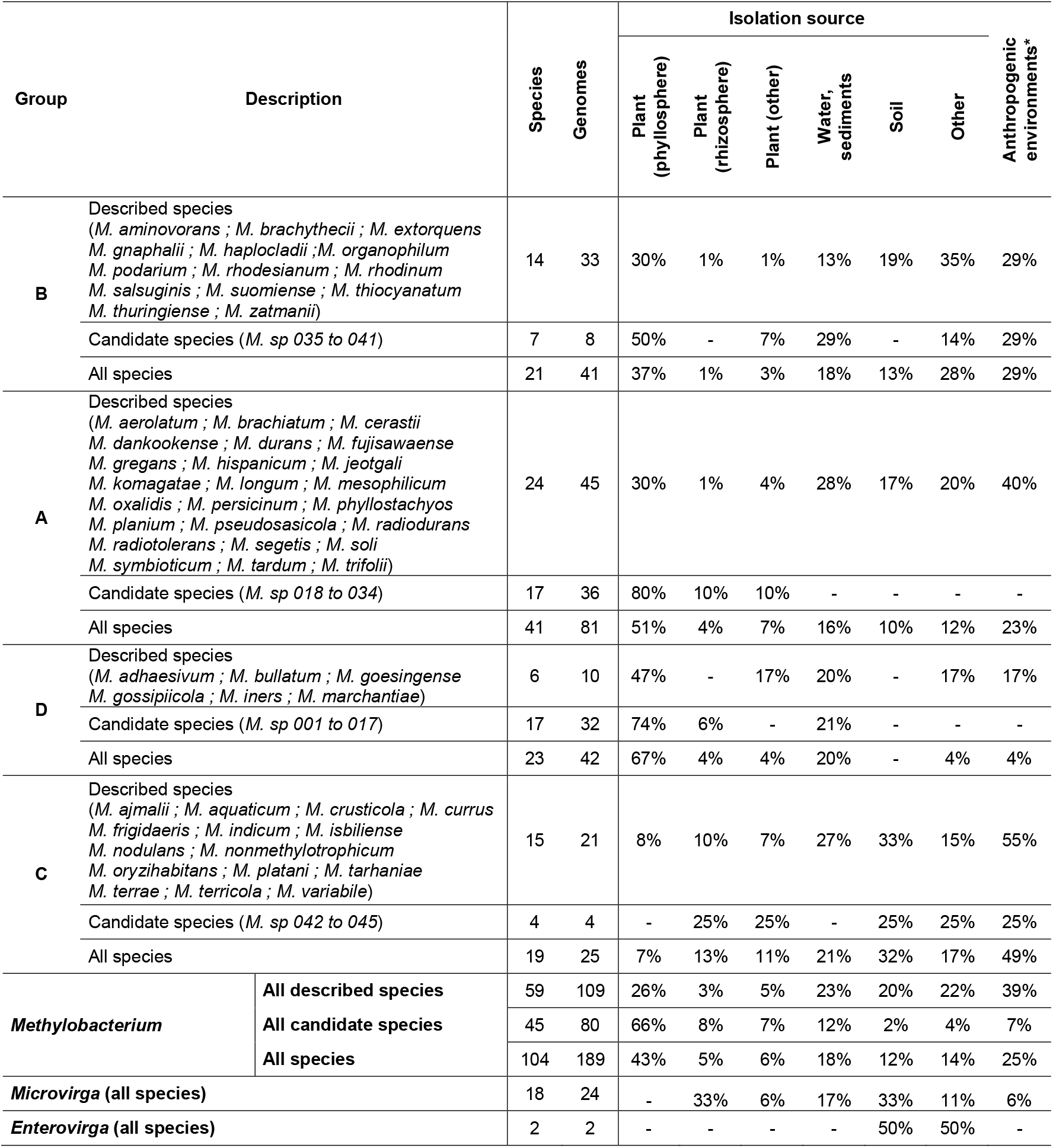
Description and isolation source of 104 *Methylobacterium* species distributed in the four main phylogenetic groups. For each group, the number of species, of genomes, and the proportion of genomes isolated from each main category of environment, are given for described and candidate species (numbered from *M. sp* 001 to *M. sp* 045), separately. Proportions were corrected by the number of genomes per species. Anthropogenic environments include several other isolations sources.

In summary, the *Methylobacterium* species are distributed across four groups, each of which with somehat distinct environmental sources of isolation (plant phyllosphere and rhizosphere, water and sediments, soils, others), as well as the proportion of strain isolated from anthropogenic environments (Table 1, Figure 1d). Group A contained 62 genomes which fell into 41 species, including 17 new species (*M. sp*. 018 to 034). Group B contained 41 genomes which fell into 21 species, including 7 candidate species (*M. sp*. 035 to 041). Group C contained 25 genomes which fell into 19 species, including 4 new candidate species (*M. sp*. 042 to 045; Table 1). Group D contained 42 genomes which fell into 23 species, including 17 new candidate species (*M. sp*. 001 to 017). Species from *Microvirga* and *Enterovirga* were mostly isolated from soil samples (65% of species; corrected by the number of genomes per species), often in association with plant roots (Rhizosphere; 30%). Species from *Methylobacterium* groups B and C were isolated from plants (40 and 31% of genomes, respectively), soil samples (13 and 32%), sediments or water samples (18 and 21%), often in association with anthropogenic environments (29 and 49%). Species from groups A and D were mostly isolated from plants (62 and 75% of species, respectively), especially the phyllosphere (51 and 67%). Of the 45 new candidate *Methylobacterium* species, most were assigned to groups A (n=17) and D (n=17); the majority (81%) was isolated from plants, and especially the phyllosphere (66%; Table 1; Figure 1d).

### *Genome comparison across* Methylobacterium *groups*

The four main *Methylobacterium* groups have consistently contrasting genome characteristics (Figure S6, Table 2). These four groups have significantly different genome sizes (Tukey test, *p*<0.001), with group D having smaller genomes (4.99 ± 0.35 Mb; Average ± SD), than groups B (5.58 ± 0.49 Mb), A (6.21 ± 0.59 Mb) and C (7.15 ± 0.66 Mb). Groups D and B had a smaller number of annotated genes (5,224 ± 476 and 5,766 ± 509, respectively) than groups A and C (6,907 ± 821 and 7,670 ± 956, respectively; *p*<0.001). The average number of gene annotation copies per genome was significantly different among groups (*p*<0.001) and was smaller for group D (1.31 ± 0.04 copies per annotation) than for group B (1.37 ± 0.05), A (1.46 ± 0.07) and C (1.54 ± 0.07). GC content was significantly lower in groups D and B (68.8 ± 1.1 and 69.1 ± 0.8 %, respectively) than in group A (70.1 ± 0.8 %; *p*<0.001) or group C (71.1 ± 0.7 %; *p*<0.001; Figure S6, Table 2). We compared the abundance of 10,187 gene annotations (excluding hypothetical proteins, repeat elements and mobile elements) across the four *Methylobacterium* groups and outgroups (Figure 2). *Methylobacterium* genomes clustered according to their gene content and abundance and matched the ASTRAL species tree (Figure 2a). As observed for other genome characteristics, group D had the smaller pan genome size (*n* = 4,217 ± 70; estimation assuming rarefaction of 15 species per group, mean and standard deviation over 100 replicates; Figure S7a), followed by group A (*n* = 4,974 ± 132), group B (*n* = 4,973 ± 137) and group D (*n* = 5,636 ± 91 genes; Figures 2b). On the contrary, group D had a larger core genome size (i.e. gene present in a single copy in all species; *n* = 1,103 ± 29 core genes) than groups A (*n*=845 ± 79), B (*n*=924 ± 65) and C (*n*=843 ± 39; Figures 2c; Figures S7b). Venn diagrams on shared annotations indicate a limited overlap of gene content among groups, with only 2,863 ± 38 pan genes shared among the four groups (Figure 2b) and 350 ± 32 core genes (Figure 2c).

**Table 2:**
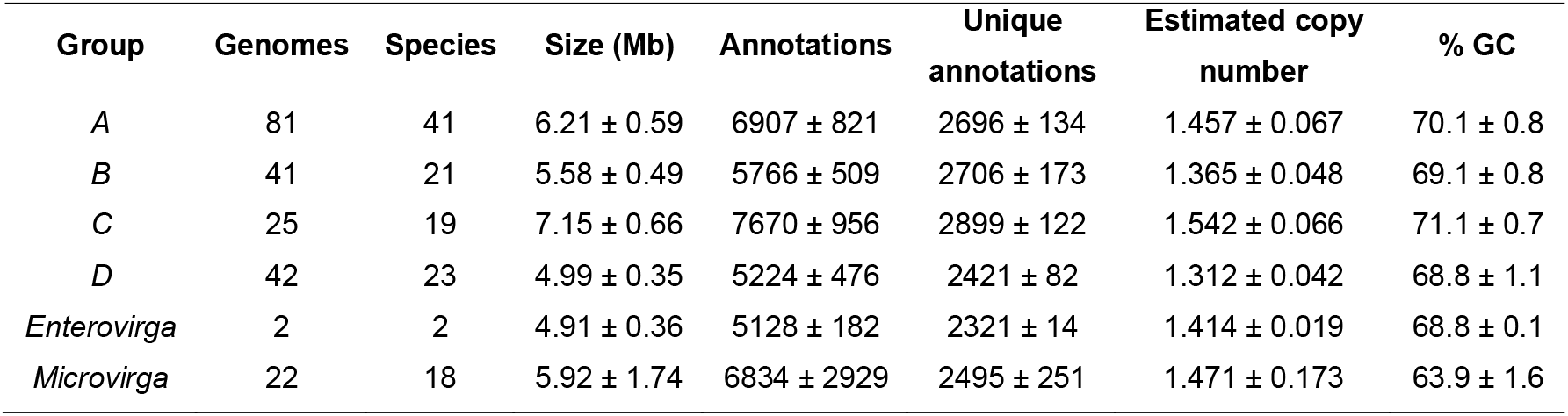
*Methylobacteriaceae* genome characteristics (average and standard deviation per group). Hypothetical protein, mobile and repeat elements were excluded from annotation counts. GC content was estimated from coding sequences.

**Figure 2:**
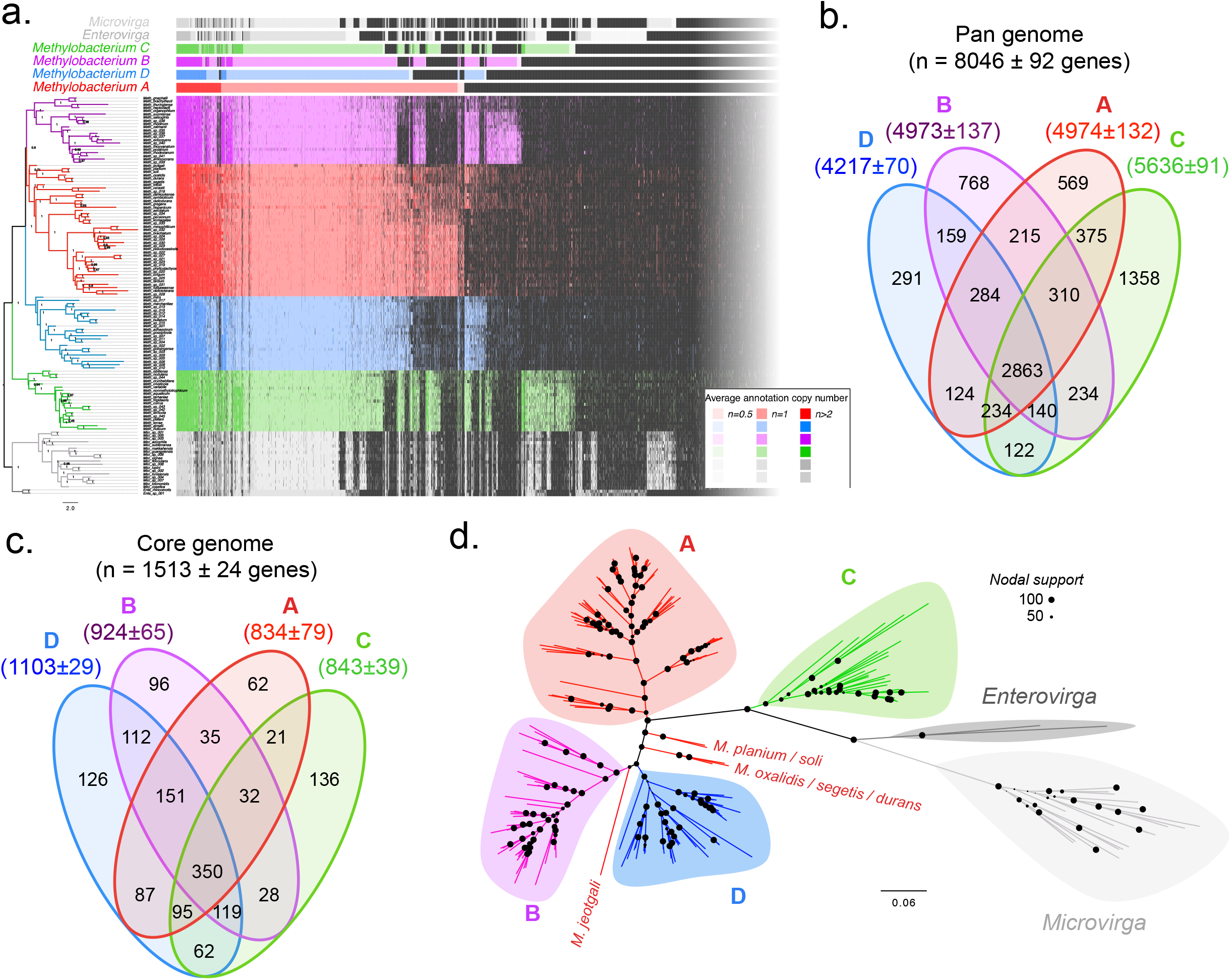
Gene content comparison among the four main *Methylobacterium* groups. a) Occurrence of 10,187 gene annotations (rows) in 124 *Methylobacteriaceae* species (average occurrence per species; column, ordered according to the ASTRAL species tree, left) and in four *Methylobacterium* groups and two outgroups (mean occurrence among species within groups; legend in bottom right) are shown. b-c) Venn diagrams showing the overlap of pan genomes (b) and core genome (c) among four groups. Pan and core genome sizes were estimated assuming 15 species per group (mean and standard deviation over 100 random resampling of 15 species per group). d) RAxML ML best tree based on annotation occurrence per genome (best ML tree, BINCAT model, 1,001 replicate trees). Main groups are shown and are monophyletic in the gene content tree, but group A: clade A2 (Alessa et al. 2021) and *M. jeotgali* branched out of group A.

### *Gene content comparison across* Methylobacterium *groups*

We asked to what extent gene content evolved concordantly along the core genome phylogeny. We used the Bray-Curtis index to measure the pairwise dissimilarity among genomes based on their gene annotation abundance (*BC*; Hellinger normalization of gene abundance; Figure 3). The dissimilarity matrix in gene content among species matched the species tree (Figure 3). Gene content was more similar among genomes from the same *Methylobacterium* species (*BC* range: 0.044 ± 0.017 - 0.080 ± 0.023) than among species within *Methylobacterium* groups (*BC* range: 0.159 ± 0.031-0.197 ± 0.044) or than among *Methylobacterium* groups (*BC* range: 0.238 ± 0.025 - 0.339 ± 0.019; Figure 3, Table 3). We determined the relationships among *Methylobacteriaceae* members upon their gene content using a ML phylogeny based on the occurrence of the 10,187 gene annotations across 213 genomes (RAxML assuming a BINCAT model; 1,001 replicate trees; Figures 2d, detailed tree in Figure S8a). The gene content tree supported each of the 124 *Methylobacteriaceae* species, as well as the monophyly of groups B, C and D (nodal support: 99, 94 and 87%, respectively). Groups A, B and D formed a monophyletic group (nodal support: 100%), making group C the most basal group, as observed for lineage (Figure 1) and species trees (Figure S5). Most of the species assigned to group A clustered together (nodal support: 77%) but five species formerly assigned to clade A2 (*M. planium, M. soli, M. oxalidis, M. durans, M. segetis*; Alessa *et al*., 2021) and *M. jeotgali* were more similar to groups B and D, which altogether formed a monophyletic group (nodal support: 97%). The normalized RF value between the gene content tree and lineage trees (Figure 1) ranged from 0.429 to 0.469. As a comparison, normalized RF values between the best gene content tree and its 1,001 replicate trees ranged from 0.085 to 0.249 (RF= 0.169±0.026), indicating that the gene content tree had significantly different topology than lineage trees.

**Figure 3:**
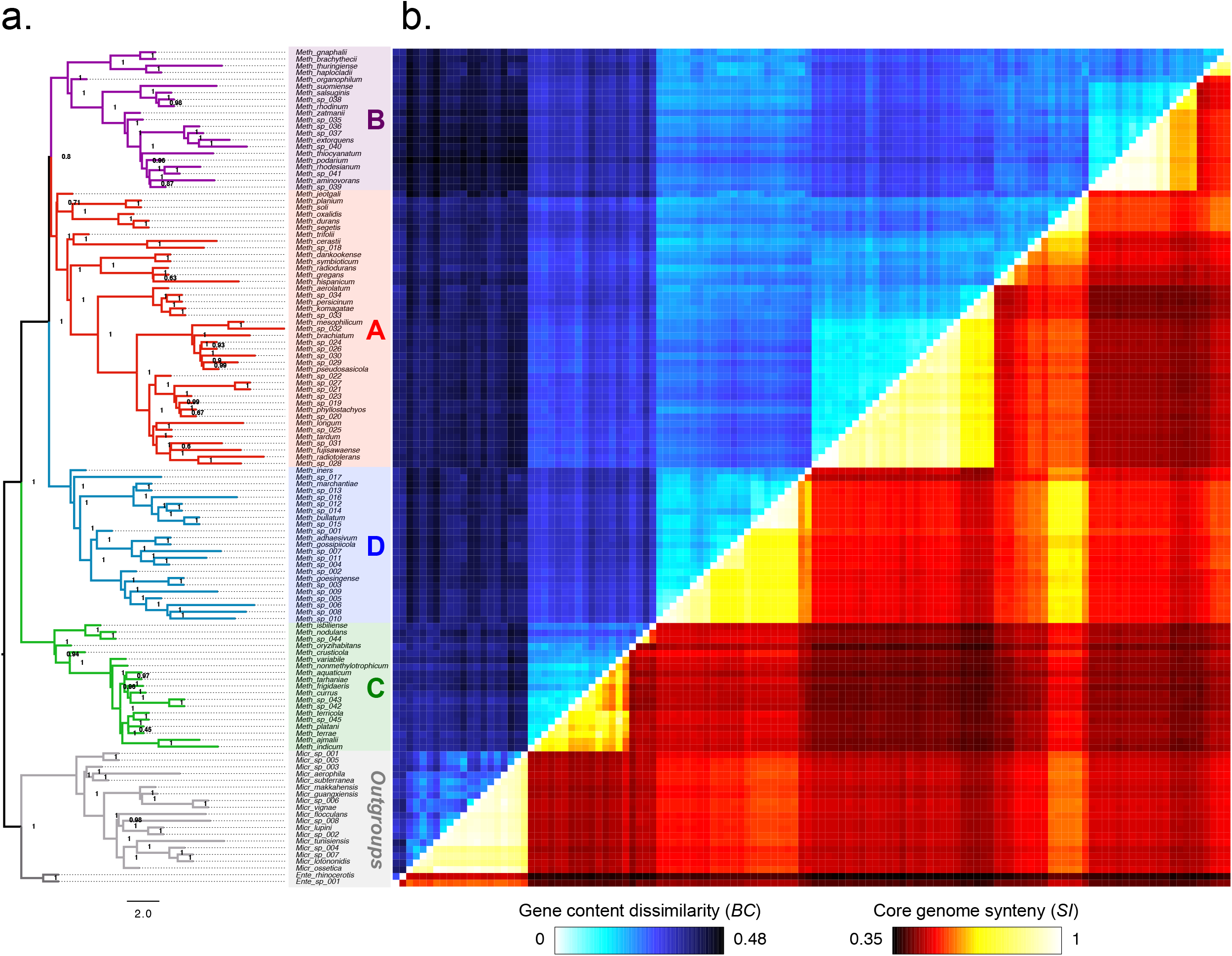
a) ASTRAL species tree. b) Heatmap of dissimilarity in gene content (*BC* index; blue scale; above diagonal) and similarity in core genome architecture (Synteny index; orange scale; below diagonal) among 104 *Methylobacterium* and 20 outgroups species, matching the species tree.

**Table 3:** Average and standard deviation in gene content dissimilarity (*BC* index, Hellinger transformation on gene occurrence per genome) per and among *Methylobacterium* group and outgroups.

| Group | *BC* within species | *BC* among species within groups | *BC* among groups |  |  |  |  |
| --- | --- | --- | --- | --- | --- | --- | --- |
|  |  |  | A | B | C | D | *Enterovirga* |
| A | 0.06 ± 0.02 | 0.19 ± 0.05 |  |  |  |  |  |
| B | 0.08 ± 0.02 | 0.17 ± 0.05 | 0.28 ± 0.02 |  |  |  |  |
| C | 0.07 ± 0.03 | 0.20 ± 0.04 | 0.29 ± 0.02 | 0.34 ± 0.02 |  |  |  |
| D | 0.04 ± 0.02 | 0.16 ± 0.03 | 0.24 ± 0.03 | 0.24 ± 0.03 | 0.32 ± 0.02 |  |  |
| *Enterovirga* | - | 0.28 | 0.39 ± 0.01 | 0.41 ± 0.02 | 0.36 ± 0.02 | 0.37 ± 0.01 |  |
| *Microvirga* | 0.03 ± 0.04 | 0.26 ± 0.04 | 0.41 ± 0.02 | 0.44 ± 0.02 | 0.38 ± 0.02 | 0.40 ± 0.02 | 0.36 ± 0.03 |

### *Core genome architecture comparison (synteny) across* Methylobacteriaceae *genomes*

We next evaluated the level of conservation in the architecture of the *Methylobacteriaceae* core genome to assess the extent of chromosomal rearrangement during *Methylobacterium* evolution. Most genomes (177 out of 213) were not fully assembled (i.e., the chromosome consisted of more than one scaffold), and we thus inferred the order or the 384 core genes along the chromosome of draft genomes by aligning their scaffolds to the chromosomes of 36 completely assembled *Methylobacteriaceae* genomes, while conserving the order of core genes within scaffolds. We compared the order of core genes among genomes using a synteny index (*SI*) calculated as the proportion of pairs of core genes (links) that were neighbors in two genomes, ranging from 0 (no link conserved) to 1 (fully conserved synteny; Figure 3, Table 4). The matrix of synteny among species was generally concordant with the species tree (Figure 3). In the 213 *Methylobacteriaceae* genomes, we observed 6,109 different links among the 384 core genes. Core genome architecture was well conserved among genomes from the same *Methylobacterium* species (*SI* range: 0.914 ± 0.064 - 0.995 ± 0.007) but was highly reshuffled among species within *Methylobacterium* groups (*SI* range: 0.608 ± 0.118 - 0.769 ± 0.207; Figure 3, Table 4). As a comparison, the core genome architecture among *Microvirga* species was remarkably well conserved (*SI* = 0.913 ± 0.048). Average synteny among *Methylobacterium* groups A, B, C and D was low (*SI* range: 0.433 ± 0.025 - 0.528 ± 0.049) and in the same order of magnitude as synteny between *Methylobacterium* and *Microvirga* genomes (*SI* range: 0.458 ± 0.010 - 0.525 ± 0.020; Figure 3, Table 4). We identified *M. planium* (strain YIM132548, group A) as the species having on average the highest core genome synteny with other *Methylobacterium* genomes. Accordingly, we used *M. planium* as a reference to visualize the conservation of the 384 links identified in its genome across *Methylobacterium* species (Figure 4a; Figure S9). We identified 150 links (involving 231 genes; 60.2% of core genes) that were mostly conserved among *Methylobacteriaceae* genomes. With the exception of a remarkably well-conserved cluster of 26 genes that included ribosomal genes and gene *rpoB* (Figure 4a; Figure S9), most of the 150 conserved links were scattered across the *M. planium* chromosome. We determined the relationships among *Methylobacteriaceae* members in their core genome architecture using a ML phylogeny based on the occurrence of 6,109 links identified across 213 genomes (RAxML assuming a BINCAT model; 1,001 replicate trees; Figure 4b, detailed tree in Figure S8b). The synteny tree supported the monophyly of the four major *Methylobacterium* groups (nodal support = 100%). Groups A, B and D formed a monophyletic group (nodal support: 83%), making group C the most basal group, as observed for lineage trees (Figure 1), species trees (Figure S5) and the gene content tree (Figure 2d). The normalized RF value between the synteny tree and lineage trees ranged from 0.589 to 0.638. As a comparison, normalized RF values between the best synteny tree and its 1,001 replicate trees ranged from 0.235 to 0.390 (RF= 0.310 ± 0.026), indicating that the synteny tree had a significantly different topology than lineage trees. Interestingly, although *M. planium* and related species previously assigned to clade A2 (*M. soli, M. oxalidis, M. segetis, M. durans*; Alessa *et al*. (2021)) as well as *M. jeotgali* and *M. trifolii* were assigned to clade A in the ML synteny tree (Figure 4b), these species had on average higher synteny with species from group D (*SI* = 0.651 ± 0.045) than with other species from group A (*SI*= 0.556 ± 0.031; Figure 4c). Accordingly, we identified 29 links involving 54 core genes that were more often conserved between groups A and D than with other *Methylobacterium* groups. These links, however, were scattered along the *M. planium* chromosome (Figures 4a, S9).

**Table 4:** Average and standard deviation in core genome synteny (*SI*) per and among *Methylobacterium* group and outgroups.

| Group | *SI* within species | *SI* among species within groups | *SI* among groups |  |  |  |  |
| --- | --- | --- | --- | --- | --- | --- | --- |
|  |  |  | A | B | C | D | *Enterovirga* |
| A | 0.98 ± 0.03 | 0.70 ± 0.17 |  |  |  |  |  |
| B | 0.99 ± 0.01 | 0.77 ± 0.21 | 0.46 ± 0.03 |  |  |  |  |
| C | 0.91 ± 0.06 | 0.61 ± 0.12 | 0.43 ± 0.03 | 0.44 ± 0.01 |  |  |  |
| D | 0.98 ± 0.02 | 0.73 ± 0.11 | 0.53 ± 0.05 | 0.51 ± 0.02 | 0.48 ± 0.01 |  |  |
| *Enterovirga* | - | 0.47 | 0.39 ± 0.03 | 0.39 ± 0.02 | 0.38 ± 0.02 | 0.42 ± 0.03 |  |
| *Microvirga* | 1.00 ± 0.00 | 0.91 ± 0.05 | 0.49 ± 0.03 | 0.48 ± 0.01 | 0.46 ± 0.01 | 0.52 ± 0.02 | 0.52 ± 0.05 |

**Figure 4:**
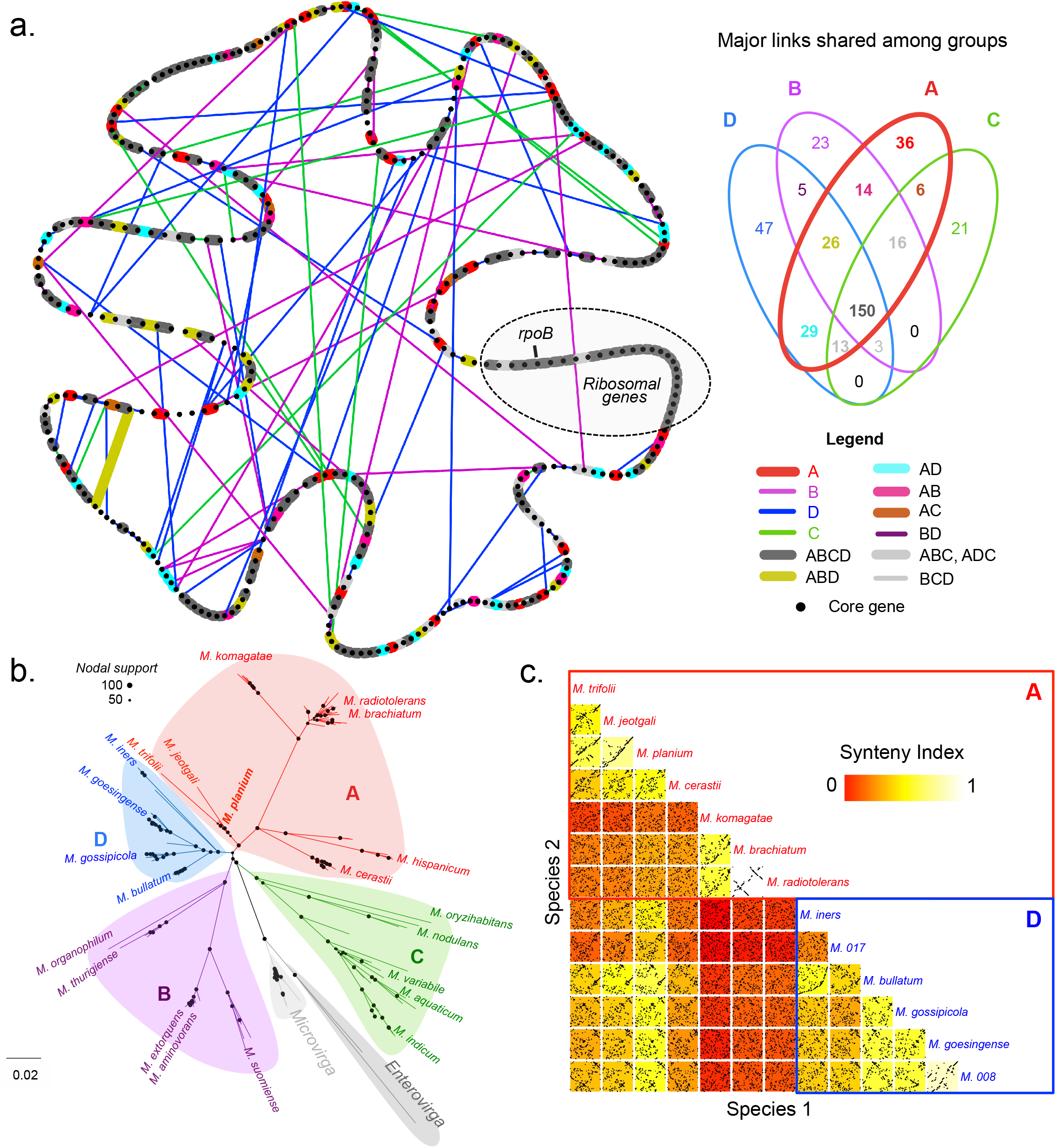
Core genome architecture comparison (synteny) among *Methylobacteriaceae* genomes. a) Consensus map of the *Methylobacterium* core genome architecture, and major rearrangements within and among *Methylobacterium* groups, using *M. planium YIM132548* core genome as a reference. The map was drawn as a network using 384 core genes as nodes, and links among neighbor core genes as edges. Only 389 links that were observed in a majority (>50%) of species from a given *Methylobacterium* group are shown (Venn diagram on top right; 5,720 links discarded). Bold lines indicate links mostly conserved in group A, colored according to their dominance in other groups (legend on bottom right). Thick lines indicate links mostly absent in group A but dominant in other groups. A syntenic island conserved in most *Methylobacterium* genomes and containing ribosomal genes and gene *rpoB* is indicated (dotted frame). b) RAxML ML best tree based on link occurrence per genome (6,109 links; best ML tree, BINCAT model, 1,001 replicate trees). Main groups are shown and are monophyletic in the synteny tree. c) Detailed synteny plot for the comparison of core genome architecture between seven species from group A and six species from group D (best assembled genome per species). For each pairwise comparison, core gene (black points) are ordered according to their relative position in species 1 genome (x-axis) and are compared with their relative positions in species 2 genome (y-axis). Each plot is colored according to the *SI* value between species 1 and 2 (scale on top right).

## Discussion

### Methylobacterium *consists of four evolutionarily divergent groups of bacteria*

Recent work has suggested that *Methylobacterium* is much more diverse than thought previously, questioning its genus status (Green and Ardley, 2018; Hördt *et al*., 2020; Alessa *et al*., 2021; Leducq *et al*., 2022). Here, we used a comprehensive phylogenomic approach to provide unprecedented insight on the taxonomic diversity of *Methylobacterium*. Our reconstructions of the *Methylobacteriaceae* lineage tree based on the core genome confirmed previous comparative genomic and phenotypic studies that group C, including *M. nodulans* and *M. aquaticum*, form a distinct and cohesive group at the root of the *Methylobacterium* phylogeny (Green and Ardley, 2018; Hördt *et al*., 2020; Alessa *et al*., 2021). On the contrary, we demonstrated that Group B, including the model species *M. extorquens*, and previously amended as a distinct genus, *Methylorubrum* (Green and Ardley, 2018), formed a monophyletic group with the *Methylobacterium* type species *M. organophilum* and other species formerly assigned to group A (e.g. *M. oxalidis* and *M. planium*) (Green and Ardley, 2018). Our analyses hence support the proposal to extend group B to *M. organophilum, M. oxalidis, M. planium*, and relatives (Alessa *et al*., 2021). Although the newly defined group B was monophyletic according to our different inferences of the *Methylobacteriaceae* species, it was still embedded within former group A (Green and Ardley, 2018), making the later paraphyletic, and confirming that *Methylorubrum* cannot be considered as a distinct genus without breaking apart *Methylobacterium* (Hördt *et al*., 2020; Alessa *et al*., 2021). Accordingly, we support the proposal to abandon “*Methylorubrum*” as a designation for group B, and to split group A into two monophyletic groups distinct from group B: group A (including *M. brachiatum, M. komagatae, M. cerastii, M. jeotgali, M. trifolii, M. planium* and relatives) and group D (including *M. bullatum, M. gossipicola, M. goesingense, M. iners* and relatives).

We observed that the newly defined monophyletic groups (A, B, C and D) were characterized by distinct genome sizes and GC content, two metrics that were highly correlated with each other in *Methylobacterium*, as observed in other bacteria (Nishida, 2012). With the exception of a few species from group A (including, *M. trifolii, M. jeotgali, M. planium* and relatives), the four groups could also be distinguished upon their gene content. GC content, genome size and gene content are widely accepted as criteria for taxonomic definition in prokaryotes (Rosselló-Mora and Amann, 2001; Coenye *et al*., 2005). We also demonstrated that core gene order was highly reshuffled among the four *Methylobacterium* groups. For instance, we observed the same level of rearrangement in core gene order among *Methylobacterium* groups, as between *Microvirga* and *Methylobacterium*, and the same level of core gene order conservation within *Methylobacterium* groups as within *Microvirga*. Core gene order has recently been proposed as a complementary criterion to define bacteria genus and species taxonomy (Chung *et al*., 2018). The fact that the four groups were monophyletic, regardless of whether we used a concatenated or a coalescent-based approach to infer the *Methylobacteriaceae* lineage tree and could be consistently distinguished from each other upon different genome characteristics (gene content, core genome architecture, GC content, genome size), supports considering them as distinct genera.

### Role of HGT and ILS in the early divergence of groups A, B and D

The evolution of bacteria is marked by recurrent HGT, gene duplication and loss events, making the reconstruction of bacterial phylogenies challenging. Given that each gene potentially has its own evolutionary history, marked by exchanges among divergent taxa, the evolutionary tree of most bacteria is quite reticulate (Shapiro *et al*., 2016). The reconstruction of a consensus phylogenetic tree (lineage tree) thus remains highly conceptual in bacteria and could only be achieved by considering a pool of genes assumed to be representative of the prevailing evolutionary history of the considered taxa: the core genome (Sakoparnig *et al*., 2021). Therefore, HGT and ILS must be considered when attempting to reconstruct bacterial phylogeny. Accordingly, we showed that the concatenated-based reconstruction of *Methylobacterium* lineage tree, assuming the same evolutionary history for each core gene, significantly differed in its topology from lineages tree reconstructions accounting for ILS and/or HGT among core genes (ASTRAL and SVDquartets lineage trees), indicating that both processes were major drivers of *Methylobacterium* evolution. While the concatenated tree suggested that groups A and D formed a monophyletic group, coalescent-based estimations from ASTRAL (ILS + HGT among genes) and SVDquartets (ILS + HGT among sites) rather indicated the earlier divergence of group D from the A/B/D group.

A possible explanation of the divergent topology in the concatenated lineage tree is that shared polymorphism was retained by ILS and/or HGT between groups A and D after the A/B divergence. Interestingly, although supporting the four *Methylobacterium* groups, our phylogeny reconstructed from core genome architecture suggested the closer relationship between groups A and D, in agreement with the concatenated lineage tree, hence supporting the hypothesis of horizontal core gene exchanges having occurred between groups A and D after the A/B divergence. Accordingly, we observed syntenic islands (groups of neighbor core genes) shared between group D and some basal species of group A (*M. jeotgali, M. trifolii, M. planium, M. oxalidis* and relatives). These islands were scattered across the *Methylobacterium* chromosome, either suggesting that extensive chromosomal rearrangements occurred after HGT between A and D, or that HGT occurred multiple times during their evolutionary history, potentially among divergent lineages. According to a phylogeny reconstructed from gene occurrence in *Methylobacterium, M. jeotgalii, M. planium, M. oxalidis* and relatives, belonged to different lineages branching at the root of groups A, B and D, supporting the hypothesis of multiple and independent gene exchanges among distinct *Methylobacterium* lineages after the divergence of the three groups, blurring their phylogenetic relationships.

### *Outstanding* Methylobacterium *diversity: the role of the phyllosphere?*

*Methylobacterium* is frequently associated with the phyllosphere, yet taxonomic and phylogenomic surveys of its diversity have mostly focused on human-impacted environments such as food factories, contaminated soils, air conditioning systems or even the International Space Station. Here we presented the first comprehensive genomic survey of *Methylobacterium* diversity in the phyllosphere. By including genomes of strains isolated from the phyllosphere of wheat (Zervas *et al*., 2019), of the model plant *A. thaliana* (Helfrich *et al*., 2018), and of trees from natural temperate forests (Leducq *et al*., 2022), our phylogenomic analysis of *Methylobacterium* revealed that its evolutionary and taxonomic diversity was larger than previously thought. In addition to recovering the 59 previously described species (Alessa *et al*., 2021), we identified 45 new (candidate) *Methylobacterium* species, of which a majority belonged to groups A and D, and were mostly isolated from the phyllosphere. Beyond taxonomic considerations, this result reveals a profound bias in our understanding of natural processes underlying the existing diversity of *Methylobacterium*, and more generally, of bacteria. For example, the evolutionary distinction between groups A and D, and their importance in *Methylobacterium* diversity, could not have been revealed without a thorough investigation of diversity in the phyllosphere, from which the majority of candidate species from groups A, B and D were isolated. A recent survey of *Methylobacterium* in metagenomes from various biomes (Lee *et al*., 2022) also suggested the association of groups B, A (represented by *M. pseudosasicola* and *M. radiotolerans* in Lee *et al*., 2022 study), and especially D (represented by *M. gossipiicola* and *M. sp*. Leaf 88) with the aerial part of plants. Similarly, we recently showed that groups A and D were the dominant *Methylobacterium* groups in the phyllosphere of trees from temperate forests (Leducq *et al*., 2022). On the contrary, groups B and C included most *Methylobacterium* model species frequently used in the lab and isolated in anthropogenic environment. While group B is occasionally identified on and isolated from the surface of leaves (Lee *et al*., 2022; Leducq *et al*., 2022), group C is rarely, if ever, found in the phyllosphere, and seems to be more widespread in soil and in aquatic environments, often in association with plant roots (Lee *et al*., 2022). Interestingly, authors from a recent study estimated that *Rhizobiales* common ancestor likely had a free-living lifestyle, while *Methylobacterium* groups A, B and D’s common ancestor likely had a plant-associated lifestyle (node 1 in Figure 1 from Wang *et al*. study (Wang *et al*., 2020)). The ancestral lifestyle of *Methylobacterium*, and more widely, of *Methylobacteriaceae*, is more unclear. The isolation source of group C genomes, as well as the two sister genera of *Methylobacterium, Enterovirga* and *Microvirga*, and their survey in metagenomes (Lee *et al*., 2022) indicate that these three groups are mostly found with soils, often in association with the rhizosphere. These observations suggest that *Methylobacteriaceae* and *Methylobacterium*’s ancestors inhabited soils, and were occasionally associated with plants, for instance in root nodules, and that *Methylobacterium* groups A/B/D’s association with the phyllosphere occurred after divergence from group C. The exact origin and nature of this association is an open question, but the smaller genome size, gene copy number and GC content we observed in group A/B/D in comparison with group C could be the genomic signatures of a progressive specialization to life on plants (Nishida, 2012).

According to our phylogenomic analyses, group D diverged first, and, like group A, was mostly isolated from the phyllosphere, suggesting that the A/B/D ancestor inhabited the surface of plant leaves. The fact that our analyses support horizontal gene exchanges between groups A, B and D is also consistent with the hypothesis that these groups lived in the same habitat during their divergence. One can speculate that some horizontally transferred genes may have had shared roles in *Methylobacterium* adaptation to the phyllosphere. For instance, strains from groups A and D were often identified in the same studies, sometimes isolated from the same plants, indicating that strains from these two groups likely share the same microhabitats on the surface of plant leaves, hence favoring gene exchanges among them and the maintenance of similar functions. Further identifications of genes exchanges among these groups and the characterization of their functions will be critical to understand evolutionary mechanisms underlying the adaptive role and radiation of *Methylobacterium* in the phyllosphere.

## Conclusion

Our unprecedented phylogenomic analysis of *Methylobacterium* revealed the outstanding diversity within this taxon, and the role of HGT in its early evolutionary history. Future genomic and functional studies will be needed to characterize the evolutionary and functional features of *Methylobacterium* adaptation to the phyllosphere. Finally, our work lays the foundation for a thorough taxonomic redefinition of this genus.

## Method

### Methylobacteriaceae *genome collection*

We assembled a collection of 213 complete and draft *Methylobacteriaceae* genomes, including 189 *Methylobacterium* and 24 genomes from related genera as outgroups (*Microvirga*: n=22; *Enterovirga*: n=2). Most *Methylobacterium* (*n*=98) and all outgroup genomes (n=24) came from distinct studies (see references in Leducq *et al*. (2022)) and corresponded to genomes publicly available in October 2020 on NCBI. We included 29 genomes from *Methylobacterium* type strains recently published (Alessa *et al*., 2021; Bijlani *et al*., 2021) in order to cover most *Methylobacterium* species described so far. We also included 38 genomes available from two large surveys of the *Arabidopsis* and wheat phyllospheres (Helfrich *et al*., 2018; Zervas *et al*., 2019) and sequenced 24 additional genomes (**see next section**) of isolates from a large survey of the temperate forest phyllosphere (Leducq *et al*., 2022), hence extending our dataset to the leaf-associated *Methylobacterium* diversity.

### *Library preparation and genome assembly of 24* Methylobacterium *strains*

We performed genome sequencing and *de novo* assembly of 24 *Methylobacterium* strains representative of the diversity previously found in the phyllosphere of two temperate forests in the province of Québec, Canada (Leducq *et al*. (2022); Dataset S2). DNA extraction was performed from culture stocks frozen at -80 °C directly after isolation and identification (Leducq *et al*., 2022) and thawed 30 min on ice. About 750 μl of cell culture were used for DNA extraction with DNeasy PowerSoil Pro Kit (Qiagen) according to the manufacturer protocol, with the following modification: final elution was repeated twice in 25 μl (total volume: 50 μl). 300 bp paired-end shotgun libraries were prepared from 35 ng genomic DNA with QIAseq FX DNA Library Kit (Qiagen) and protocol was adjusted to target DNA fragments in the range 400-1000 bp. Genomes were assembled from libraries with MEGAHIT (Li *et al*., 2015) with default parameters. Genome assemblies had 7050-24785 contigs with average depth in the range 188-304x and a total size in the range 7.2-17.1 Mb. After removing contigs with depth <10x, we obtained 82-411 contigs per genome. Most assemblies had total size (5-7 Mb) and average GC content (67-70%) in the expected range for *Methylobacterium* genomes (Dataset S2). For three out of twenty-four genomes, GC content and depth distribution were clearly bimodal, and total size was much higher (9.5-11.9 Mb) suggesting that these assemblies contained genomes from at least two evolutionary distinct taxa. For these three heterogeneous assemblies, we divided contigs into two pools based on median depth value between two modes (threshold range: 100-150x). For each heterogeneous assembly, the pool with highest average depth (174-241x) had average GC content (67-68%) and total size (5.6-5.8 Mb) in the ranges expected for *Methylobacterium*. Contigs with lower depth were considered as contaminants and discarded from assemblies.

### Gene annotation

*Methylobacteriaceae* genomes (n=213) were individually annotated using RAST (https://rast.nmpdr.org/rast.cgi) (Aziz *et al*., 2008) with following parameters: genetic code=11; Annotation scheme=RASTtk; Preserve gene calls=no; Automatically fix errors=yes; Fix frameshifts=no; Backfill gaps=yes. Annotation output from each genome was retrieved separately as Spreadsheet (GFF file in tab-separated text format). Core genome definition was conducted in R (R-Developement-Core-Team, 2011). For each genome, we retrieved the abundance of gene annotations (column *function* in RAST output), excluding *Hypothetical proteins, repeat regions* and *Mobile element proteins* (Dataset S3).

### Methylobacteriaceae *core genome definition*

We first defined the *Methylobacteriaceae* core genome from 184 genomes, excluding 29 genomes that were not yet published nor annotated at the time of the analysis (Alessa *et al*., 2021; Bijlani *et al*., 2021). In these 184 genomes, we identified 9,970 unique gene annotations (i.e., regardless copy number: Dataset S3), with on average 2637 ± 210 unique gene annotations per genome. We defined candidate core genes as genes present in one copy in at least 90% of the 184 genomes, resulting in 893 candidate core genes, for which we retrieve the nucleotide sequence (column *nucleotide_sequence* in RAST output). In order to correct for false gene duplication events that increased consistently with assembly incompleteness (Figure S1) and to estimate the actual copy number of each candidate core gene, we used 36 complete *Methylobacteriaceae* genomes as references (defined as genomes with *N50* > 3.10^6^ Mb). For each candidate core gene, we calculated the average expected nucleotide sequence size observed among 36 complete genomes. Then, for each genome (n=184) and each candidate core gene, we retrieved all nucleotide sequences (0-10 per gene and genome) and calculated their average size normalized (divided) by the average nucleotide sequence size observed in complete genomes. By doing so, we could distinguish between duplication caused by genome incompleteness (single copy genes divided between different scaffolds) and real duplication events (Figure S2). We considered 398 genes for which at least one genome had more than one copy with normalized size >0.75 as true duplicates and removed them from candidate core genes. For the 495 remaining candidate core genes, we considered single-copy genes with normalized size >1.3 and gene copies with normalized size <0.7 (regardless copy number) as missing data in the considered genome. After this filter, we removed 111 candidate core genes that were missing in at least 4 genomes, resulting in 384 core genes for which a single full-length copy could be retrieved for at least 181 genomes (out of 184; Dataset S4). Subsequently, we included recently sequenced genomes from 29 *Methylobacterium* species type strains that were missing from our survey (Alessa *et al*., 2021; Bijlani *et al*., 2021). By doing this, we slightly extended the number of unique gene annotations in *Methylobacteriaceae* (*n* = 10,190). We confirmed that the 384 previously identified core genes were part of the *Methylobacteriaceae* core genome and retrieved each core gene nucleotide sequence for at least 26 out of 29 genomes. Our final dataset consisted in 213 genomes for which we retrieved 327 to 384 core genes nucleotide sequences (381 ± 6; mean, SD; Dataset S1).

### Core gene nucleotide sequence alignments

We performed an alignment for each core gene. For each genome (n = 184 + 29 = 213), we first extracted nucleotide sequences of the 384 core genes (when not missing data for the considered genome; column *nucleotide_sequence* in RAST output) and converted them in sequence fasta files using R package *seqinr()*. We then performed an alignment for each core gene using R packages *seqinr()* and *msa()*. For each gene, nucleotide sequences were translated (function *getTrans()*) and alignments of amino-acid sequences were performed using ClustalW with default parameters in function *msa()*. Sequences were converted back in nucleotides (stop codons excluded) and 5’ and 3’ end codons with more than 90% of missing data (gaps of “Ns”) were trimmed. We also constructed an alignment of concatenated core genes nucleotide sequence alignments. In the concatenated alignment, sequences of genes missing for at least one of the 213 genomes (0-6 genomes missing per gene) were replaced by strings of “Ns”.

### *Inferences of the* Methylobacteriaceae *lineage trees*

We reconstructed the lineage tree of *Methylobacteriaceae* from 213 genomes from the 384 core gene nucleotide sequences using three complementary approaches in order to assess the effect of ILS and HGT in the evolutionary history of *Methylobacterium*.

First, we used RAxML v. 8.2.8 (Stamatakis, 2014) to determine a maximum-likelihood (ML) lineage tree from concatenated alignments of the core 384 gene nucleotide sequences assuming a different substitution model for each gene but the same evolutionary tree for all gene (and hence not accounting for ILS or HGT). First, we used PartitionFinder implemented in IQ-tree2 (Minh *et al*., 2020, p.) to determine an appropriate bipartitioning scheme allowing us to merge genes evolving under similar nucleotide substitution models (Lanfear *et al*., 2012). The best-fit partition scheme was determined using TESTMERGERONLY model (option *–m*) to avoid tree reconstruction, and using the relaxed hierarchical clustering algorithm to reduce the computation burden (Lanfear *et al*., 2014) by only examining the top 10% partition merging schemes (option *–rcluster*). Second, we inferred the *Methylobacteriaceae* lineage tree from the 384 core gene alignment with RAxML v. 8.2.8 (Stamatakis, 2014), using the IQ-tree2 best-scheme output file as partition file (option –*q* in RAxML). The program performed 512 replicate (bootstrap) searches from independent starting trees with a GTRCAT model of substitution, estimating parameters for each partition separately. Of the 512 trees, the one with the highest ML score (the best-scoring tree) was retained as the lineage tree.

Second, we used ASTRAL-III (Zhang *et al*., 2018), a coalescent-based method inferring the lineage and the species trees by combining individual core gene trees, hence accounting for ILS and HGT among genes. For each core gene, a gene tree was first inferred from nucleotide sequence alignments with RAxML v. 8.2.8 (Stamatakis, 2014). Briefly, for each gene, the program performed 1,000 replicate (bootstrap) searches from independent starting trees assuming a GTRgamma model of nucleotide substitution. Each gene tree in Newick format, including branch length (*L*: nucleotide substitution per site) and node label (*N*: nodal support representing the proportion of replicated supporting nodes), was imported in R as a vector. The gene tree in RAxML format: *((():L1[N1]):L2[N2])* was rewritten so that it could be readable in R (package *ape* (Paradis and Schliep, 2019)) and ASTRAL-III: *((():L1)N1:L2)N2*. The tree was then reopened in R with function read.tree (package *ape*) and nodes with < 10% support were collapsed using function *collapseUnsupportedEdges* (package *ips*), to optimize accuracy in estimating the lineage and species tree (Zhang *et al*., 2018). All reformatted gene trees were written in a single file (*multiPhylo* object), which was used to infer the lineage and the species tree in ASTRAL-III v5.7.7, with default parameters. In ASTRAL trees, branch lengths were measured in coalescent units and nodal support represented local posterior probability (Sayyari and Mirarab, 2016).

Third, we used SVDquartets (Chifman and Kubatko, 2014) as implemented in PAUP* v4.0a (build 169) (Wilgenbusch and Swofford, 2003), a coalescent-based method estimating the tree for each possible combination of four genomes and assuming all sites unlinked in the concatenated alignment of 384 genes. This accounts for both ILS and HGT within and among genes.

We estimated both species trees and lineage trees from the concatenated 213 *Methylobacteriaceae* core genes by evaluating 2,000,000 random quartets for 100 bootstrap replicates. Phylogenies were estimated under the multispecies coalescent model accounting for incomplete lineage sorting (ILS) and assessing all sites independently to account for recombination within and among loci.

Lineage trees were displayed in Figtree v1.4.4 and rooted on *Microvirga* and *Enterovirga*.

### Methylobacteriaceae *species definition and lineage tree inferences*

We classified *Methylobacteriaceae* genomes in species using percentage nucleotide similarity (*PNS*) on the core genome (concatenated alignments on 384 core genes; 361,403 bp), similar to average nucleotide identity (Mende *et al*., 2013; Chun and Rainey, 2014). Briefly, *PNS* between two genomes was calculated in R as the proportion of conserved nucleotide positions, gaps and “Ns” excluded (Dataset S4). Two genomes were considered from the same species when their *PNS* was higher or equal to 97%, a threshold similar to what is typically used for bacterial species (96.5% based on nucleotides sequences of 40 marker genes; (Mende *et al*., 2013; Chun and Rainey, 2014)). We inferred the *Methylobacteriaceae* species tree using both ASTRAL-III (Zhang *et al*., 2018) and SVDquartets (Chifman and Kubatko, 2014), as described above, using individual assignment to species determined from PNS. Species trees were displayed in Figtree v1.4.4 and rooted on *Microvirga* and *Enterovirga*.

### Test for HGT and ILS severity in lineage and species tree inferences

We tested for the severity of HGT and ILS in our dataset by measuring the differences in tree topologies estimated using different assumptions. To quantify differences between the topologies obtained under different assumptions, we calculated normalized Robinson-Foulds (RF) distances (Robinson and Foulds, 1981), which evaluates the pairwise proportion of unique nodes between tree topologies, between all three lineage trees (RAxML from concatenated core gene alignments, SVDquartets, and ASTRAL) as well as between both species trees (ASTRAL and SVDquartets). RF distances were estimated using the treedist function implemented within PAUP* v4.0 (build 169) (Wilgenbusch and Swofford, 2003) using final phylogenies in NEWICK format as input. We also calculated the distribution of RF distances between our best RAxML tree from concatenated core gene alignments and all 512 RAxML bootstrap replicates. We then compared our RF distances between each inference method to this distribution of distances to assess whether discordant topologies among lineage trees are due to different assumptions of methods, or due to phylogenetic uncertainty.

### Methylobacteriaceae *genome characteristics and gene content*

We analyzed *Methylobacteriaceae* characteristics (size, GC content and gene content) of the coding genome. For each genome, we first calculated the number of gene annotations, their total nucleotide size (coding genome size), their GC content, the number of unique annotations (excluding hypothetical proteins, repeat elements and mobile elements), and the average copy number of annotations (Dataset S1, Table 2). For each statistic, we compared *Methylobacterium* groups (as defined by lineage trees) and outgroups (*Microvirga, Enterovirga*) using a Tukey test.

In a heatmap, we displayed the average abundance of 10,187 gene annotations (excluding hypothetical proteins, repeat elements and mobile elements) per *Methylobacteriaceae* species, per *Methylobacterium* group and outgroup, ordered according to the ASTRAL species tree (Figure S5a). For gene abundance per species, we calculated the average occurrence (*n*) of each gene annotation across genomes assigned to the same species, rounded to 0 (*n*<0.5), 1 (0.5≤*n*<1.5) or 2 copies (*n*≥2). For gene abundance per group, we calculated the average occurrence of each gene annotation across species assigned to the same group, using the same principle as for species.

We estimated pan genome and core genome sizes per *Methylobacterium* group (unknown proteins, repeat and mobile elements excluded; Figure S7). To deal with biases in size estimations due to the variable number of genomes per group, we applied rarefaction on gene number estimates by randomly sampling 1 to *N* genomes per group (Park *et al*., 2019) and by forcing resampling of one genome per species. For each *N* value and each group, we calculated the average and standard deviation in core genome size (genes in 1:1 copy in each genome of a given group) and in pan genome (any gene present in at least one copy in at least one genome of a given group) over 100 replicates. As pan genome size estimations increased with the number of sampled species per group (Figure S7a) and core genome size estimations decreased (Figure S7b), curves of estimates per group did not cross each other, nor reached a plateau, indicating that sizes were either under-estimated (pan genomes) or over-estimated (core genome) but could still be compared among groups. Accordingly, we compared pan and core genome sizes among *Methylobacterium* groups in a Venn diagram, assuming 15 species per group (Figures 2b,c,).

We constructed a phylogeny of *Methylobacteriaceae* based on gene content (Figure 2d). First, we constructed a matrix of gene occurrence in each genome (0 for absence and 1 for presence) and converted it into a fasta file (one sequence per genome). We inferred an evolutionary tree of based on gene content using with RAxML v. 8.2.8 (Stamatakis, 2014). The program performed 1,000 replicate (bootstrap) searches from independent starting trees with a BINCAT model of substitution assuming gene presence of absence as binary data. Of the 1,000 replicate trees, the one with the highest Maximum-likelihood (ML) score (the best-scoring tree) was considered as the best tree. The tree was displayed in Figtree v1.4.4 and rooted on *Microvirga* and *Enterovirga*.

In order to compare *Methylobacteriaceae* genomes based upon their gene content, we calculated an index of dissimilarity among each pair of genomes from their gene abundance (Table 3, Figure 3b). As no index was available for this purpose, we used the Bray-Curtis (*BC*) index of dissimilarity, initially developed in ecology for the comparison of communities based on their species abundance (Bray and Curtis, 1957). To minimize the effect of higher copy number due to false gene duplications due to genome incompleteness, we applied a normalization on gene abundances (Hellinger normalization; function *decostand* in R package *vegan*). We calculated pairwise *BC* indexes of dissimilarity among normalized gene abundances, using function *vegdist* in R package *vegan*).

### Core genome architecture (synteny)

We evaluated the level of conservation in core gene organization (synteny) between *Methylobacteriaceae* genomes. All analyses were performed in R (R-Developement-Core-Team, 2011). For each genome, we retrieved core gene coordinates (scaffold name and coordinates in the scaffold). For complete genomes consisting in a single linear scaffold (n=36), each core gene was paired with its two immediate neighbors, based on shortest distance between gene *start* and *stop* coordinates, and core genes located on scaffold edges were also paired together, assuming genome circularity. Hence, for each complete genome, each of the 384 core genes was involved in two links (pairs of neighbor core genes), for a total of *N*=384 links per genome. For the 177 draft genomes, core genes were located on different scaffolds, so we predicted scaffold order and orientation using complete genomes as references. The draft genome with the highest completeness was reorganized first. Briefly, for each comparison with a reference genome, and for each scaffold of the draft genome, the list of embedded core genes was retrieved, and a score based on gene average order in the reference genome was calculated. Scaffolds of the draft genome were reordered according to these scores (one per scaffold). Then, each scaffold was eventually reoriented (without affecting gene order within scaffolds) to optimize pairing of edge genes (genes located at the edge of a scaffold) as compared to the reference genome. We repeated the operation for comparisons with the 36 reference genomes. Finally, for each of the 36 new configurations of the draft genome, we calculated a synteny conservation index (*SI*) with each reference genome, as the proportion of links conserved between two genomes. *SI* ranged from 0 (no link conserved) to 1 (fully conserved synteny). The draft genome configuration with the highest *SI* value was conserved for further analyses and added to the list of reference genomes. We repeated this operation for each draft genome, ranked according their decreasing completeness, hence increasing the number of references genomes and possible configurations for highly fragmented genomes. Finally, we calculated *SI* for all possible pairwise comparison between genomes (Figure 3b), and average and standard deviation values within and among *Methylobacteriaceae* species and groups (Table 4).

In order to visualize the spatial organization of core genes along the *Methylobacteriaceae* chromosome, we used two approaches. First, we realized a heatmap of link conservation per species, along a reference genome (Figure S9). We choose as reference the genome having the highest average *SI* value with other *Methylobacterium* genomes. In the heatmap, we displayed the 384 links identified in the reference genome, ordered according to core gene order along its chromosome, and highlighted them when also present in other *Methylobacteriaceae* species. For each species, we also reported the average *SI* value with the reference genome. Finally, for each link in the reference genome, we calculated its frequency in each *Methylobacterium* group. Second, we drawn a consensus map of the *Methylobacterium* core genome architecture, as well as major rearrangements within and among *Methylobacterium* groups, as a network in Cytoscape v.3.4.0 (Shannon *et al*., 2003) (Figure 4a). In this network, we represented the 384 core genes as nodes, ordered according to *M. planium YIM132548* core genome, and links among neighbor core genes as edges. The network was drawn using 389 links observed in a majority (>50%) of species from a given *Methylobacterium* group (5,720 links discarded).

In order to reconstruct the evolution of *Methylobacteriaceae* core genome based on its architecture, we constructed a matrix of occurrence of each possible link observed among genomes (0 for absence and 1 for presence) and converted it into a fasta file. We inferred an ML evolutionary tree of *Methylobacteriaceae* based on synteny using with RAxML v. 8.2.8 (Stamatakis, 2014) with a BINCAT model of substitution assuming pair of core genes presence of absence as binary data, as described for annotations (Figure 4b).

## Data availability statement

Draft genomes for 24 *Methylobacterium* strains corresponding to Bioproject PRJNA730554 (Biosamples listed in Dataset S1) were deposited on NCBI under accession numbers JAKSXU000000000 - JAKSYR000000000. R code and related data were deposited on Github (https://github.com/JBLED/methylobacterium-phylogenomics.git).

## Acknowledgments

This research received funding from the National Science Foundation to C.J.M., N.C.M.-G., J.A.F, J.M.S., and J.A.L (DEB-1831838).

We thank Benjamin Oswald and the IBEST Computational Resources Core at University of Idaho for their technical support, and Alex Alleman, as well as xxx anonymous reviewers for their helpful suggestions on the manuscript.

J.-B. L., D. S., M. S. and J.M.S. performed the bioinformatic analyses, J.-B. L., D.C.-M., and G.B. performed the experiments, N. C. M.-G., J. A. L., J. A. F. and S. S. provided discussion at the early stage of the study, J.-B. L., B. J. S., S. W. K., J. M. S. and C. J. M. drafted the manuscript, with help from N. C. M.-G., J. A. L. and S. S.

