## Supplementary material for "Comprehensive phylogenomics of *Methylobacterium* reveals four evolutionary distinct groups and underappreciated phyllosphere diversity": Figure S1

**Figure S1** : Proportion of genes per genome present in 1 (blue), 2 (green), 3 (orange), 4 (red) or 5 (purple) copies per genome in function of genome assembly quality (given by the number of scaffolds per genome, log scale).

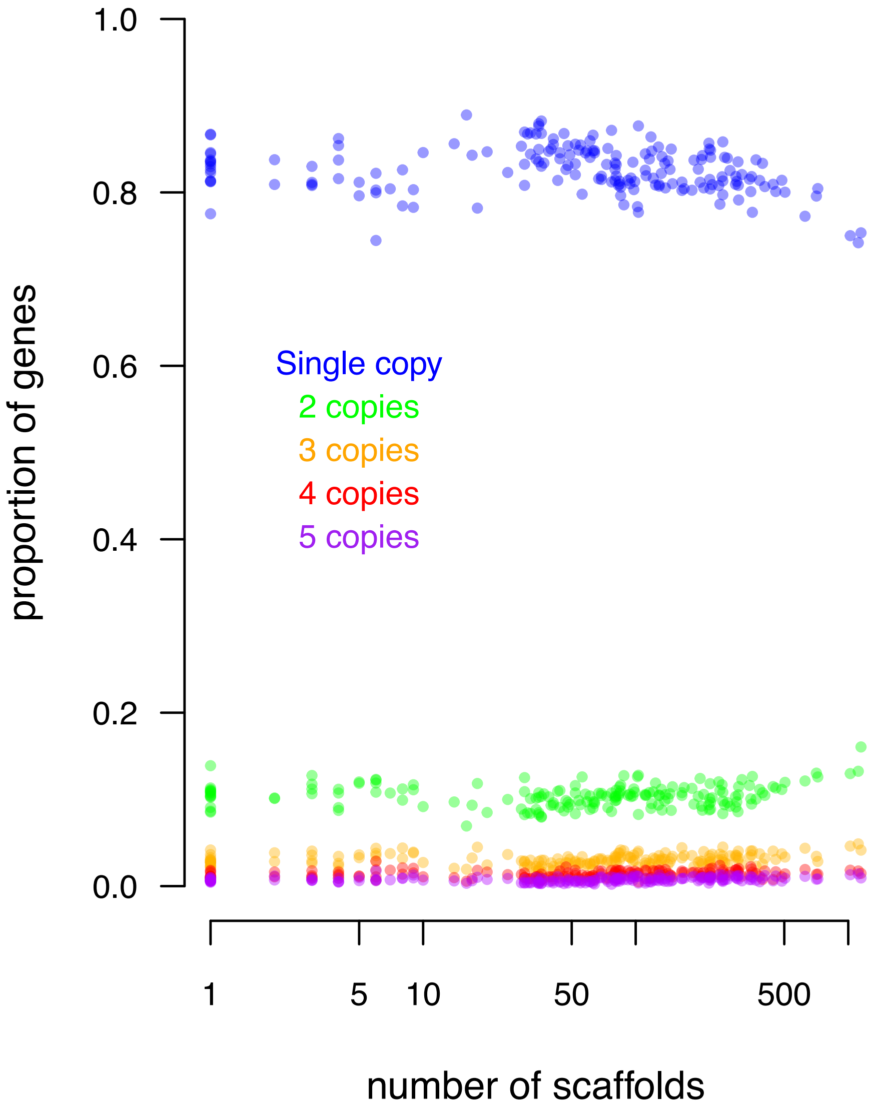
