## Supplementary material for "Comprehensive phylogenomics of *Methylobacterium* reveals four evolutionary distinct groups and underappreciated phyllosphere diversity": Figure S2

**Figure S2:** Identification of true core genes among 893 candidate core genes present in a single copy in at least 90% of 184 *Methylobacteriaceae* genomes. Average gene size normalized (divided) by the average nucleotide sequence size observed in complete genomes (defined as genomes with *N50* > $3.10^6$ Mb) was plotted against the number of copies observed per genome. Each dot represents one copy in one genome. Lines represent the expected copy number for each normalized size/observed copy number combination. 398 genes for which at least one genome had more than one copy with normalized size >0.75 were considered as true duplicates and removed from the analysis (red). For the 495 remaining candidate core genes, single-copy genes with normalized size >1.3 and gene copies with normalized size <0.7 (regardless copy number) were considered as missing data (blue). Of the remaining genes, 384 genes with a single copy in at least 180 genomes were considered as true core genes.

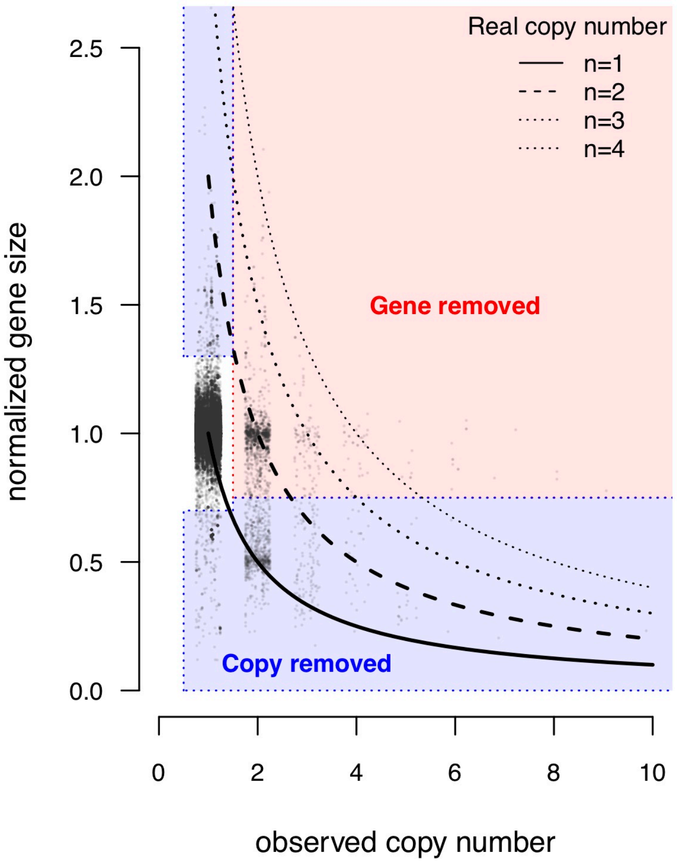
