## Supplementary material for "Comprehensive phylogenomics of *Methylobacterium* reveals four evolutionary distinct groups and underappreciated phyllosphere diversity": Figure S3

**Figure S3:** Detailed lineage trees of *Methylobacteriaceae* reconstructed from 213 genomes using ASTRAL and SVDquartet. a) ASTRAL tree inferred from 384 core gene ML trees. Each gene ML tree was inferred in RAxML assuming a GTRgamma model (1,000 replicated trees; nodes with less than 10% of support collapsed) and combined in ASTRAL-III. Branch lengths are in coalescent units. Nodal support values represent local posterior probability. b) SVD quartet tree inferred from the concatenated alignments of 384 core gene nucleotide sequences. Nodal support values indicate the proportion of quartets supporting each node. Trees were rooted on *Microvirga* and *Enterovirga*. Branches and genomes names (Genus, species, strain) were colored according to assignation to *Methylobacterium* groups (A: red; B: purple; C: green; D: blue) and outgroups (*Microvirga*: grey; *Enterovirga*: dark grey)

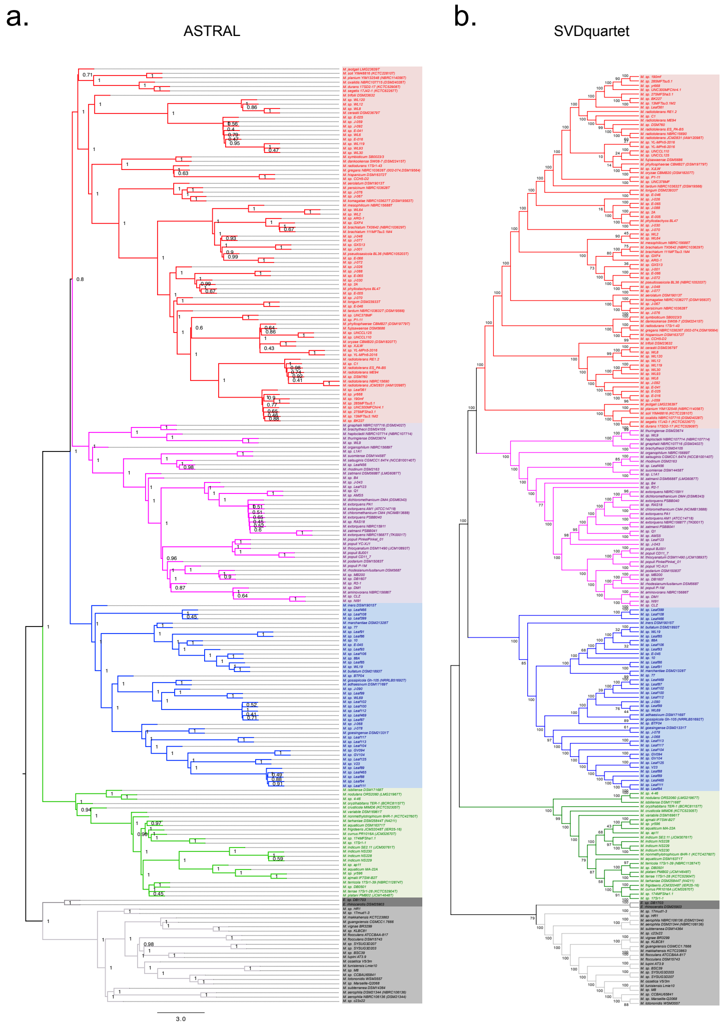
