## Supplementary material for "Comprehensive phylogenomics of *Methylobacterium* reveals four evolutionary distinct groups and underappreciated phyllosphere diversity": Figure S4

**Figure S4:** Normalized RF distance distribution between the RAxML majority consensus rule lineage tree and the 512 replicate trees (grey; see Figure 1a) and normalized RF distances between lineage trees (points; legend on top right).

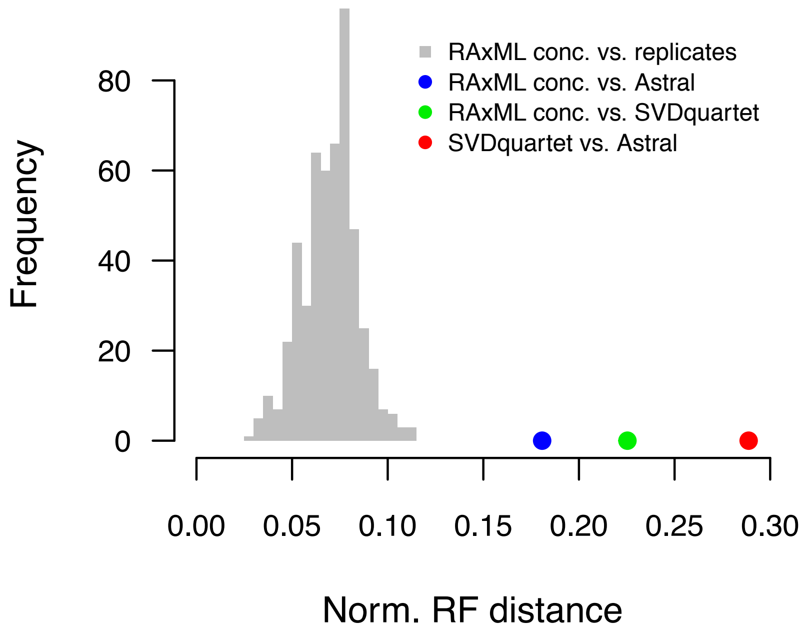
