## Supplementary material for "Comprehensive phylogenomics of *Methylobacterium* reveals four evolutionary distinct groups and underappreciated phyllosphere diversity": Figure S6

**Figure S6:** Comparison of genome characteristics among 4 *Methylobacterium* groups. Genome size, number of unique gene annotations per genome (unknown proteins, repeat elements and mobile elements excluded), average gene copy number per genome, and GC content (in coding genome) are compared to each other. Each point represents values for a genome, colored according to assignment to *Methylobacterium* groups (A: red; B: purple; C: green; D: blue). Ellipses indicate 50% of standard deviation centered on average values for each group.

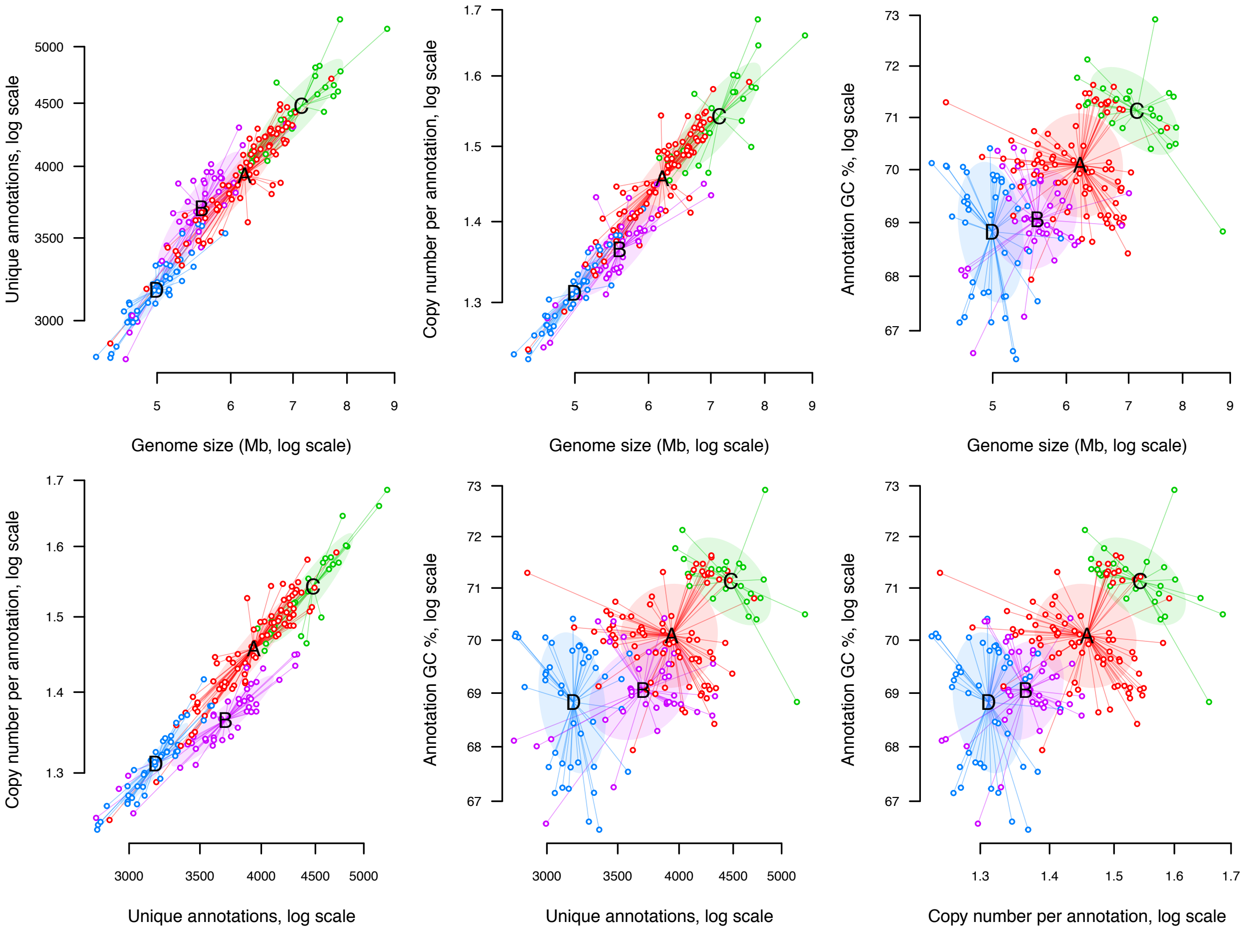
