## Supplementary material for "Comprehensive phylogenomics of *Methylobacterium* reveals four evolutionary distinct groups and underappreciated phyllosphere diversity": Figure S7

**Figure S7:** Pan (left) and core (right) genome size estimations in four *Methylobacterium* groups and *Microvirga*. Genome sizes per group (number of genes per groups; y-axis) were estimated for every number of species assumed in the range 1-n (n = maximum number of species per group; x-axis). For each group and number of species, average (lines) and standard deviation (frames) over 100 random resampling of n species per group were estimated. Dotted lines indicate the value for which pan genome and core genome size where estimated (n=15 species per group; Figure 2).

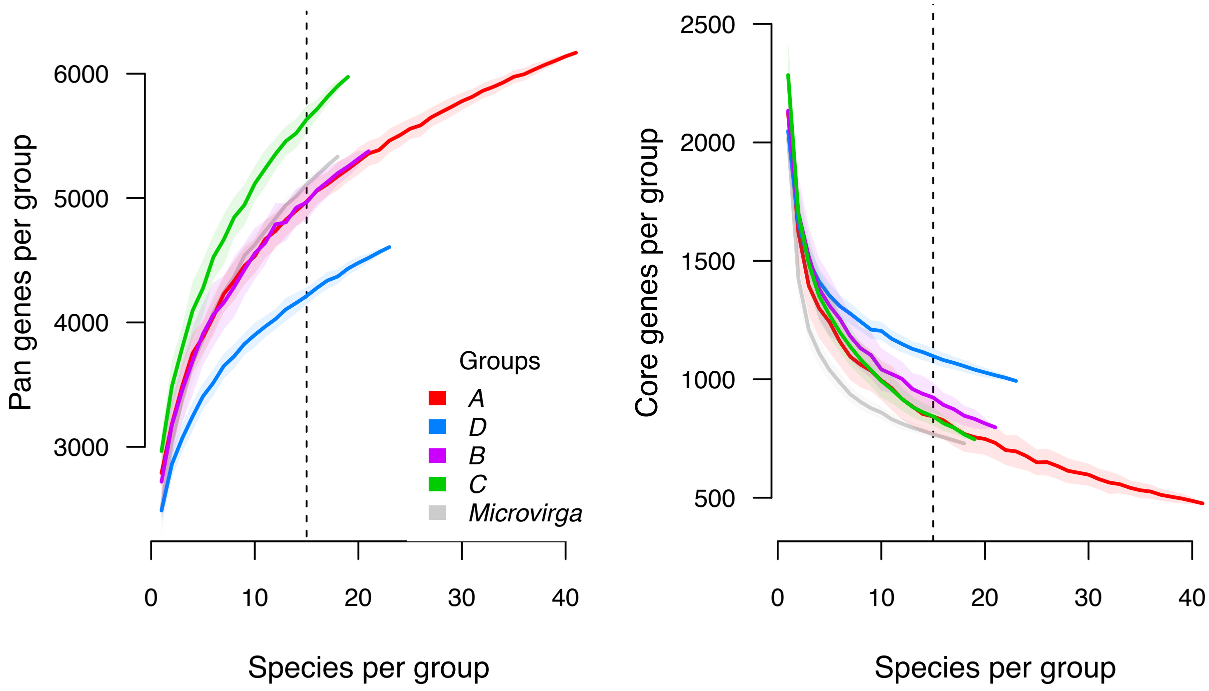
