## Supplementary material for "Comprehensive phylogenomics of *Methylobacterium* reveals four evolutionary distinct groups and underappreciated phyllosphere diversity": Figure S8

**Figure S8:** Detailed *Methylobacteriaceae* lineage trees inferred from gene content (a) and core genome synteny (b). Each ML tree was inferred in RAxML assuming a BINCAT model (1,000 replicated trees). Nodal support values indicate the proportion of replicate tree supporting each node. Trees were rooted on *Microvirga* and *Enterovirga*. Branches were colored according to assignation to *Methylobacterium* groups (A: red; B: purple; C: green; D: blue) and outgroups (*Microvirga*: grey; *Enterovirga*: dark grey)

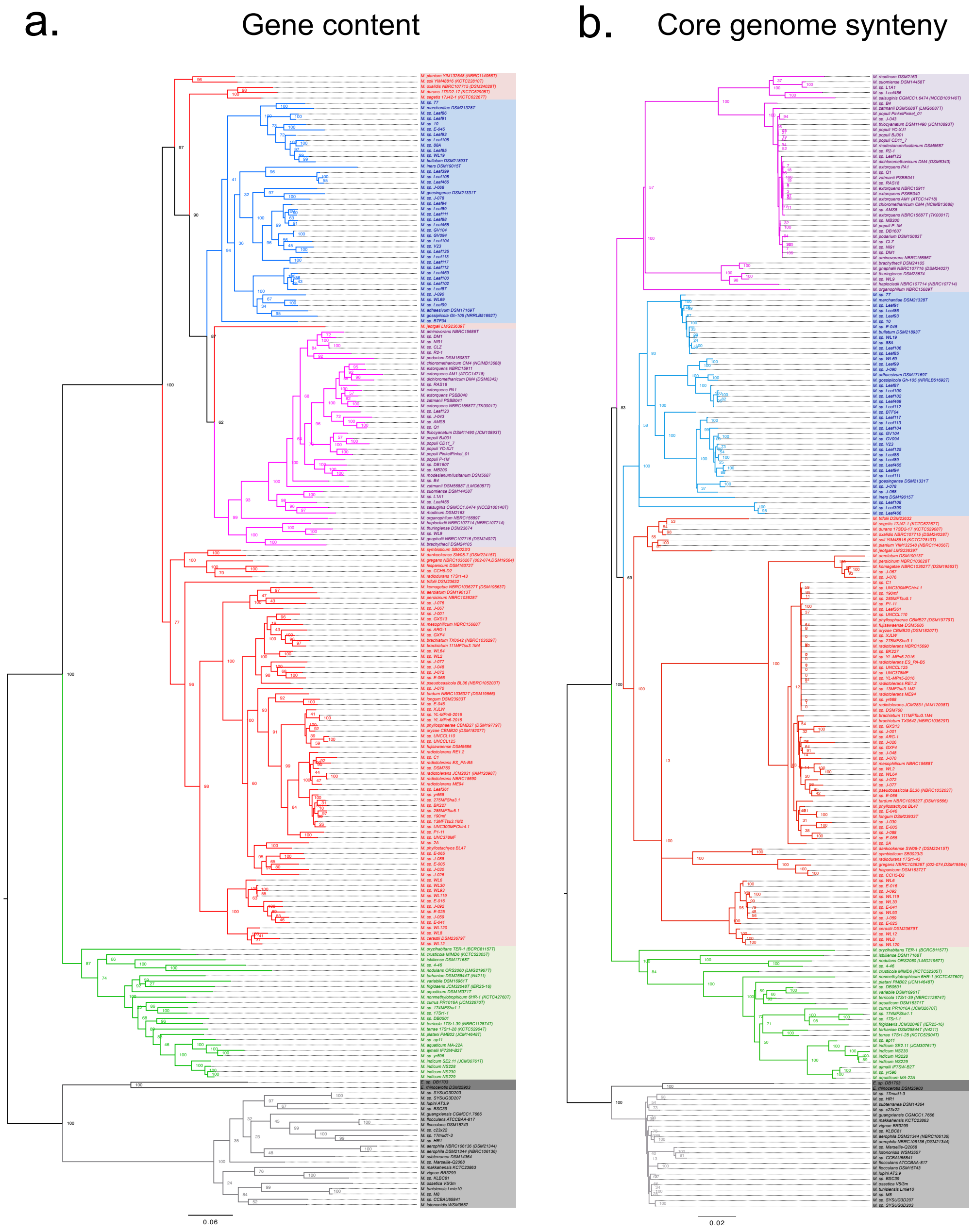
