## Supplementary material for "Comprehensive phylogenomics of *Methylobacterium* reveals four evolutionary distinct groups and underappreciated phyllosphere diversity": Figure S9

**Figure S9:** Comparison of core genome architecture among 124 *Methylobacteriaceae* species (rows; ordered according to the ASTRAL species tree, left) using 384 links (pairs of contiguous core genes) observed in the *M. planium* genome as a reference (links are ordered according to the reference genome). For each species, *M. planium* links are colored according to their group (B: purple; A: red; D: blue; C: green; outgroups: grey) when observed. Average *SI* values per link per *Methylobacterium* group are indicated in the top diagram. Blue points indicate links that are more conserved in group D than group A. Average *SI* values with *M. planium* are indicated per species on the right diagram.

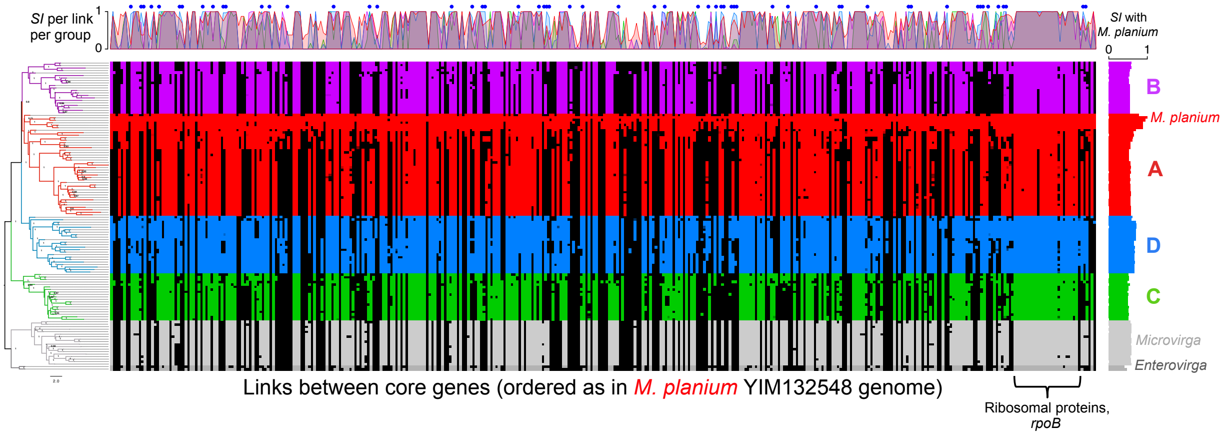
